# Transcriptomic and proteomic regulation through abundant, dynamic, and independent arginine methylation by Type I and Type II PRMTs

**DOI:** 10.1101/2020.06.23.167601

**Authors:** Stephanie M. Lehman, Hongshan Chen, Emmanuel S. Burgos, Maxim Maron, Sitaram Gayatri, Edward Nieves, Dina L. Bai, Simone Sidoli, Varun Gupta, Matthew R. Marunde, James R. Bone, Zu-Wen Sun, Mark T. Bedford, Jeffrey Shabanowitz, Donald F. Hunt, David Shechter

## Abstract

Arginine methylation is essential for both cellular viability and development and is also dysregulated in cancer. PRMTs catalyze the post translational monomethylation (Rme1/MMA, catalyzed by Type I-III), asymmetric (Rme2a/ADMA, Type I enzymes)-, or symmetric (Rme2s/SDMA, Type II enzymes) dimethylation of arginine. Despite many studies, a thorough integration of PRMT enzyme substrate determination and proteomic and transcriptomic consequences of inhibiting Type I and II PRMTs is lacking. To characterize cellular substrates for Type I (Rme2a) and Type II (Rme2s) PRMTs, human A549 lung adenocarcinoma cells were treated with either Type I (MS023) or Type II (GSK591) inhibitors. Using total proteome hydrolysis, we developed a new mass spectrometry approach to analyze total arginine and lysine content. We showed that Rme1 was a minor population (∼0.1% of total arginine), Rme2a was highly abundant (∼1.1%), and Rme2s was intermediate (∼0.4%). While Rme2s was mostly eliminated by GSK591 treatment, total Rme1 and Rme2a were more resistant to perturbation. To quantitatively characterize substrate preferences of the major enzymes PRMT1, PRMT4(CARM1), and PRMT5, we used oriented peptide array libraries (OPAL) in methyltransferase assays. We demonstrated that while PRMT5 tolerates aspartic acid residues in the substrate, PRMT1 does not. Importantly, PRMT4 methylated previously uncharacterized hydrophobic motifs. To integrate our studies, we performed PTMScan on PRMT-inhibited A549 cells and enriched for methylated arginine containing tryptic peptides. For detection of highly charged peptides, a method to analyze the samples using electron transfer dissociation was developed. Proteomic analysis revealed distinct methylated species enriched in nuclear function, RNA-binding, intrinsically disordered domains, and liquid-liquid phase separation. Parallel studies with proteomics and RNA-Seq revealed distinct, but ontologically overlapping, consequences to PRMT inhibition. Overall, we demonstrate a wider PRMT substrate diversity and methylarginine functional consequence than previously shown.

## Introduction

Methylation, a conserved post-translational modification (PTM), is the enzymatic transfer of a methyl group by a methyltransferase from the donor, S-adeno-syl-L-methionine (SAM), to a substrate amino acid. Protein methylation primarily occurs on lysines and arginines (Levy 2019; Lorton and Shechter 2019). While arginine methylation is estimated to be fairly abundant and it is known to have significant roles in transcription, RNA splicing, and DNA repair–especially in cancer (Guccione and Richard 2019)–where it is found and what its functional cellular consequences are still incompletely defined. Indeed, there is a disparity in understanding the overall abundance of this PTM, with experiments indicating 0.1% to 7% of the total proteome arginine methylated (Paik and Kim 1970; Boffa et al. 1977; Bulau et al. 2006; Dhar et al. 2013; Larsen et al. 2016).

Vertebrates have nine <u>p</u>rotein arginine <u>m</u>ethyltrans-ferases (PRMTs), grouped into three types (Guccione and Richard 2019; Tewary et al. 2019). Types I, II, and III PRMTs mono-methylate the terminal nitrogens of the arginine guanidinium group (ω-NG-monomethylarginine) (Rme1 or MMA). Type I PRMTs (PRMT1,2,3,4,6,9 [PRMT4 is also known as CARM1]) further modify the mono-methylated nitrogen to produce an asymmetrically di-methylated arginine residue (Rme2a or ADMA) (ω-NG,NG-asymmetric dimethylarginine). Type II PRMTs (5 and 9) further methylate the unmodified guanidinium nitrogen, thereby creating a symmetrically dimethylated arginine residue (Rme2s or SDMA) (ω-NG,NG’-symmetric dimethylarginine). PRMT5 is the predominant Type II enzyme (Stopa et al. 2015). Importantly, while PRMT1 exhibits some substrate rebinding (Gui et al. 2013), PRMT enzymes are generally distributive (Wang et al. 2013; Wang et al. 2014; Burgos et al. 2015) and scavenge each other’s substrates (Dhar et al. 2013). These observations suggest a complex enzyme/substrate interplay and imply potentially complementary cellular mechanisms. Both PRMT1 and PRMT5–considered the primary enzymes for Rme2a and Rme2s, respectively–are essential for cell viability and development (reviewed by Guccione and Richard 2019).

Many of the known roles for arginine methylation in the cell involve protein binding or DNA and RNA binding (Lorton and Shechter 2019). Studies on histone arginine methylation highlight the importance of the PTM’s geometry. Both histone H3R2 and H4R3 are either symmetrically or asymmetrically modified (by at least PRMT1, PRMT4, and PRMT5) leading to inverse cellular transcriptional consequences: H3R2me1, H3R2me2s, and H4R3me2a are “activation” marks, while H3R2me2a and H4R3me2s are “repressive” marks. For instance, we and others have shown that the WDR5 reader structurally distinguishes between the various methylarginines (Hyllus et al. 2007; Migliori et al. 2012; Chen et al. 2017; Lorton et al. 2020).

Hydrogen binding, water displacement, and electrostatic interactions are the mechanism of arginine:RNA binding (Hofweber and Dormann 2019). The methylation pattern can direct binding preferences: both the number of methylations and the symmetry status are relevant for function (Lorton and Shechter 2019). Therefore, understanding how many proteins, at what sites, and what kind of methylation is found is critical for future mechanistic studies.

To determine the total methylarginine content of cells, others have used protein hydrolysis and HPLC chromatography (Paik and Kim 1970; Boffa et al. 1977; Bulau et al. 2006; Dhar et al. 2013). To characterize PRMT enzymes’ substrate motifs, a variety of targeted enzymatic and computational approaches have been used (Wooderchak et al. 2008; Gathiaka et al. 2016; Gayatri et al. 2016). Finally, others have tested the proteomic and transcriptomic consequences of inhibition of various PRMTs (Chan-Penebre et al. 2015; Eram et al. 2016; Fong et al. 2019; Musiani et al. 2019; Radzisheuskaya et al. 2019).

To characterize the arginine methylome, recent investigations have used a variety of approaches. For instance, PTMScan–previously developed to immuno-precipitate post-translationally modified peptides after lysate protease cleavage (Stokes et al. 2012)–was used in HEK293 cells to identify thousands of PRMT-regulated Rme1 sites (Larsen et al. 2016) and to identify PRMT1 substrates in *Toxoplasma* (Yakubu et al. 2017). Others have used PTMScan with SILAC to identify sites of all three methylarginine states, mostly found to be enriched in RNA-binding proteins and regulating splicing (Fong et al. 2019; Musiani et al. 2019; Radzisheuskaya et al. 2019). Middle-down proteomics coupled with electron-transfer dissociation (ETD) was used to specifically identify arginine methylation in arginine- and serine-rich domains in RNA-binding proteins (Kundinger et al. 2020).

Substrate motifs for PRMTs have also been explored: most studies show that glycine and arginine rich regions, or GAR motifs are common targets for arginine methylation (Branscombe et al. 2001). Prior studies revealed that CARM1 prefers PGM motifs (proline, glycine, and methionine rich sequences)(Cheng et al. 2007; Gayatri et al. 2016; Shishkova et al. 2017). PRMT7 methylates an ‘RxR’ motif (Feng et al. 2013).

We set out to provide a larger and complete picture of the arginine methylome, PRMT substrate motifs, specific methylated proteins, and the PRMT-regulated proteome and transcriptome. Therefore, to elucidate the diverse cellular consequences of the family of PRMTs1-9, here we develop and integrate many different techniques, including: a new total amino acid mass spectrometry analysis; PRMT enzyme peptide substrate array library assays; PTMScan using electron-transfer dissociation (ETD) for all three methylarginine states; proteomics, and transcriptomics. We demonstrate how drugging either type I or type II PRMTs lead to independent consequences. Overall, we provide a new and broader view of the role of PRMTs in cellular metabolism.

## Results

### Total proteome arginine methylation

To determine the role of Type I and Type II PRMTs in the catalysis of total proteome arginine methylation, we treated human A549 lung adenocarcinoma cells with DMSO (control), 1 µM GSK591 (PRMT5 inhibitor), or 1 µM MS023 (Type I PRMT inhibitor) for one week (**Figure 1a**). Neither treatment resulted in substantial change in any of the PRMT enzyme protein expression (**Supplemental Figure S1a**) but they did result in expected histone H4R3 methylation changes (**Supplemental Figure S1b**). Using 8M Urea, we prepared total cell lysates, precipitated total protein, and hydrolyzed the proteins to their constituent amino acids (**Figure 1b**). We developed UHPLC methods to separate the four arginine species: R, Rme1, Rme2a, and Rme2s (**Figure 1c**). Using standards and triple quadrupole (QqQ) mass spectrometry, we developed methodology to characterize each arginine isoform fragment ion loss (**Figure 1d**). An example MS/MS spectrum showing identification and quantification of Rme2s is shown in **Figure 1e**. In each of the three treated cell states, we then used this analysis to quantify total proteome Rme1, Rme2s, and Rme2a as a percentage of total arginine. A standard curve of each arginine species response in our method–using NMR determined concentrations–is shown in **Supplemental Figure S2a-d**. As shown in **Figure 1f**, monomethylarginine (Rme1) constituted roughly 0.1% of the total arginine and surprisingly was not appreciably affected by either GSK591 or MS023 treatment. Symmetric dimethylarginine (Rme2s) constituted rougly 0.4% of total arginine and, as expected, the vast majority was lost by GSK591 treatment. Rme2a constituted 1.1% of the total cellular arginine and roughly 40% of this was lost by MS023 treatment. We also developed similar approaches for lysine (K, Kme1, Kme2, Kme3); the methyllysine content in untreated cells was only about 0.6% of total lysine (**Supplemental Figure S2e**).

**Figure 1.**
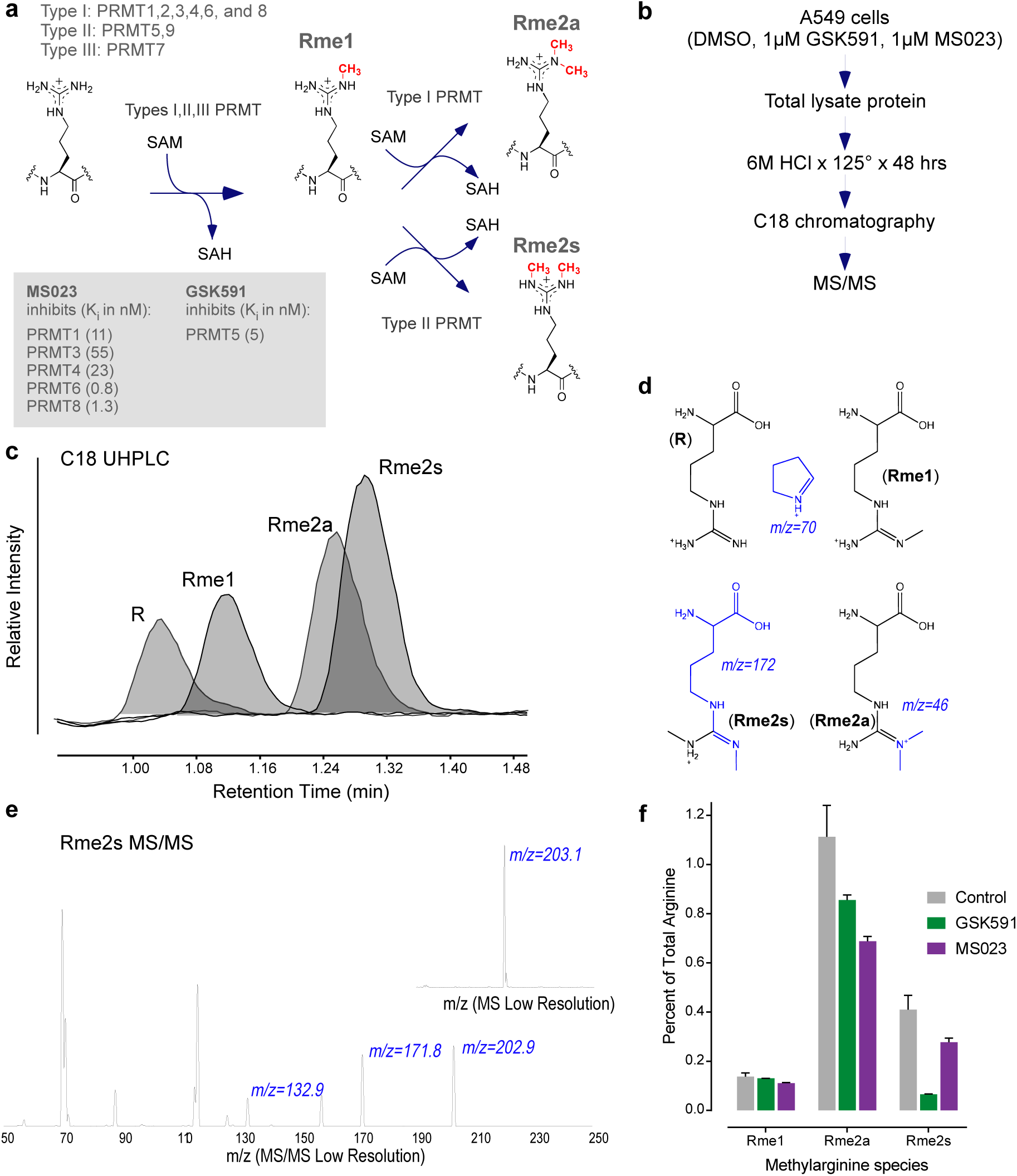
Total proteome arginine methylation. **a.** Schematic of the reactions catalyzed by the three types of protein arginine methyltransferases (Type I PRMT catalyze Rme1 and Rme2a; Type II PRMTs catalyze Rme1 and Rme2s; Type III PRMTs catalyze only Rme1) **b.** Experimental setup: A549 cells were cultured for 1 week with either DMSO, 1 µM GSK591, or 1 µM MS023. Total protein lysates (8M Urea) were precipitated with TCA followed by complete hydrolysis in 6M HCl and heat. Resulting amino acid products were separated on C18 reversed-phase chromatography and subjected to MS/MS analysis. **c.** UHPLC trace of the four Arginine amino-acids resolved onto Agilent Zorbax Eclipse Plus C18 RRHD column (2.1×150 mm, 1.8 µm; kept at 40 °C). **d.** Representation of Arginine amino-acid species detected within the study (R, Rme1, Rme2a and Rme2s). The fragments detected and utilized as quantifiers within the low resolution set-up are depicted in blue; likewise, their matching m/z are annotated **e.** As an example, the fragmentation pattern (MS/MS trace) of the Rme2s species from its parent peak (insert; m/z = 203.1). **f.** Calculated abundance of each methylarginine species as a percent of total arginine are shown (DMSO control in gray, GSK591-treated cells in green, and MS023-treated cells are shown in purple).

### Substrate specificity of PRMTs

To gain further information about PRMT substrates and characterize the enzymatic specificities of the major and most abundant enzymes (PRMT1, PRMT4/CARM1, and PRMT5), we used oriented peptide array libraries (OPAL) and recombinant enzymes. OPAL substrate peptides contained a fixed arginine (R) surrounded by one of any of 19 amino acids (cysteine was not included) in fixed positions, with the remainder as a degenerate mix of amino acids (**Figure 2a**). We then, in the presence of ^3^H-SAM, incubated recombinant human PRMT1, human PRMT4 (CARM1), *C. elegans* PRMT5, and *Xenopus laevis* PRMT5-MEP50 complex on the OPAL arrays. Using scintillation counting on the OPAL peptides, we determined relative methyltransferase activity and for each enzyme plotted the results as a heatmap (**Figure 2b-e**)(Cornett et al. 2018). To determine consensus motifs, we generated Weblogo probability plots (**Figure 2f-i, bottom panels**). We observed a few striking features of the methylated motifs: PRMT1 methylated “GR”-rich substrates and accommodated some hydrophobic residues but did not methylate substrates containing acidic residues; PRMT4/CARM1 methylated the known “PR”-rich motifs with additional enrichment of “F/W”-rich motifs, did not prefer glycine or acidic residues, and overall strongly preferred hydrophobic residues; both the *C. elegans* PRMT5 that is active without WDR77/MEP50 and the vertebrate PRMT5-MEP50 complex methylated “GR”-rich substrates and also accommodated acidic aspartic acid residues. These results revealed previously unknown differences in *in vitro* PRMT substrate specificity.

**Figure 2.**
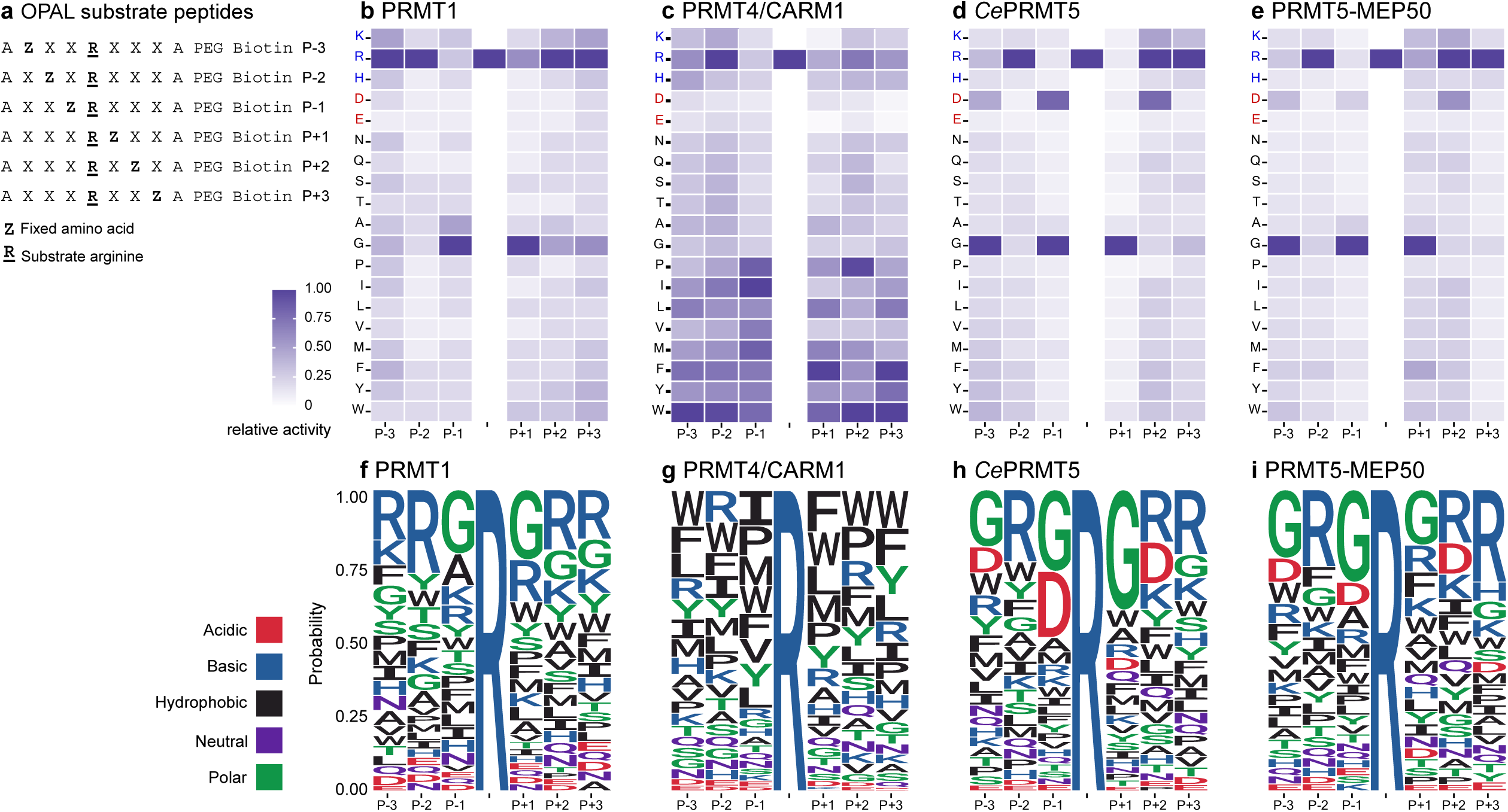
In vitro substrate specificities of the major enzymes PRMT1, PRMT4/CARM1, and PRMT5. **a.** Oriented peptide array library (OPAL) substrate degeneracy schematic. As shown, the substrate arginine (R) is fixed as is one other position per mixture (Z) in positions (−3,-2,-1, or 1,2,3 relative to the fixed R). The remainder of the substrate peptide residues are degenerate (Z). **b.** Homo sapiens (Hs) PRMT1 relative activity towards the OPAL substrate library. Each row represents that fixed amino acid in each position. Charged residues are colored (blue = positive, red = negative) and shown at the top. Relative activity is shown as a heatmap (0 to 100%, white to blue). **c.** HsPRMT4/CARM1 relative activity, as in b. **d.** C. elegans (Ce)PRMT5 relative activity, as in b. **e.** HsPRMT5-MEP50 complex relative activity, as in b. **f.** Sequence logo probability plot of PRMT1 relative activity. Acidic residues are shown in red, basic in blue, hydrophobic in black, neutral in purple, and polar residues shown in green. **g.** Sequence logo probability plot of *Hs*PRMT4/CARM1 relative activity. **h.** Sequence logo probability plot of *Ce*PRMT5 relative activity **i.** Sequence logo probability plot of *Xl*PRMT5-MEP50 relative activity

### PTMScan identification of methylated proteins

Thus far, our studies revealed that methylarginine levels are higher than expected and that the family of PRMT enzymes has a more diverse substrate specificity than previously shown. To characterize which proteins are methylated by the family of PRMTs, we turned to the PTMScan approach. We first probed A549 cell total lysate–as treated in each condition–with the Cell Signaling Technology (CST) Rme1, Rme2s, and Rme2a antibodies (**Figure 3a**) (Gayatri et al. 2016). Importantly, as shown, these antibodies are highly biased towards substrates containing multiple GR-rich methylated epitopes. Considering our initial total methylarginine analysis and *in vitro* enzymology assays, this methodology is therefore highly biased towards a subset of PRMT substrates.

**Figure 3.**
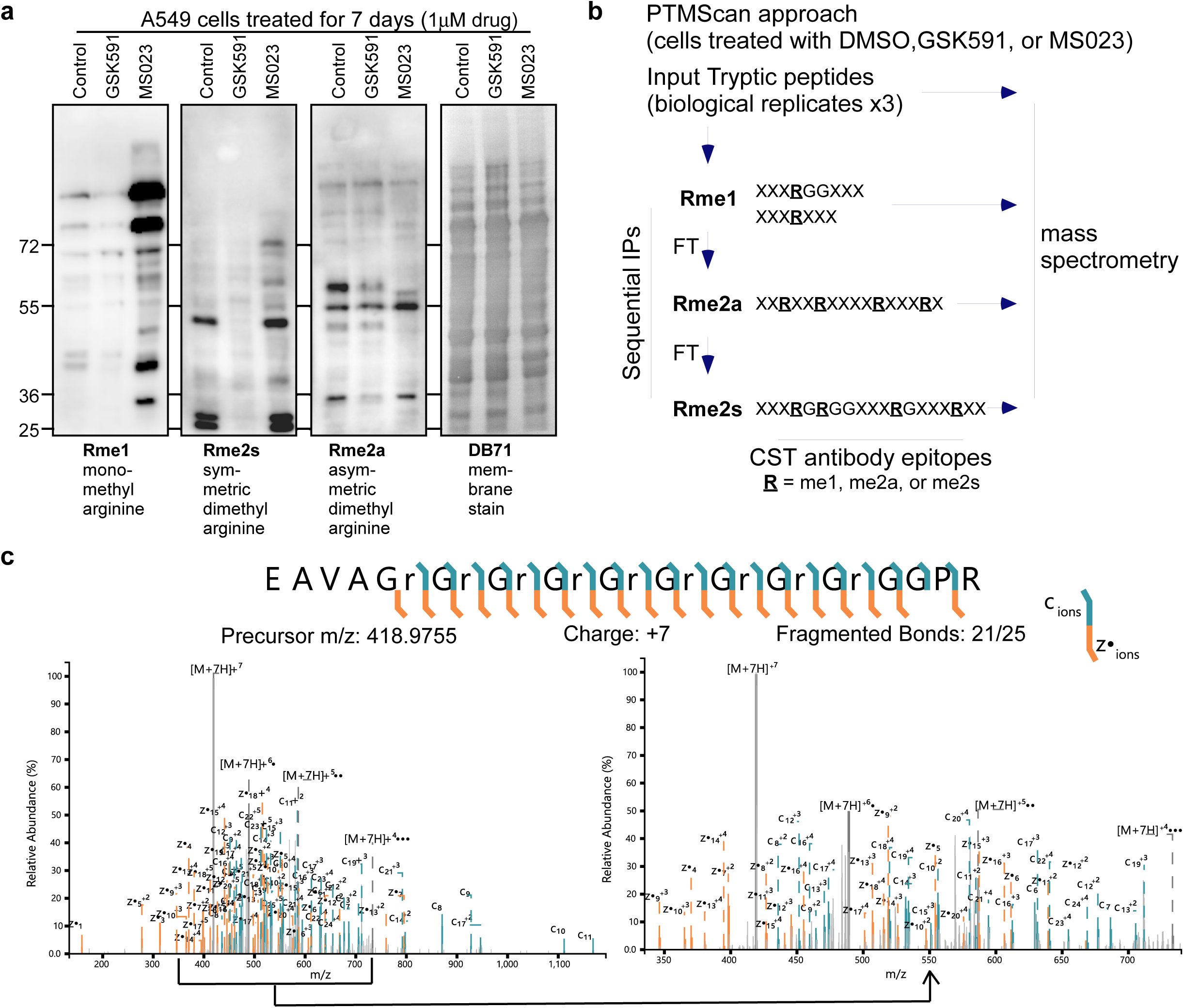
PTMScan of arginine methylated proteins in cells treated with PRMT inhibitors. **a.** Total proteome western blots of all three methylarginine states are shown. Using the CST Rme1 (left), Rme2s (center left panel), and Rme2a (center right panel) antibodies, the changes in methylarginine protein abundance is shown for each of the control (DMSO), GSK591, and MS023 conditions. The right panel shows the Direct Blue 71 (DB71) membrane stain total loaded protein. **b.** Schematic of PTMScan approach. Purified tryptic peptides, in biological triplicate, were successively immunoprecipitated with the Rme1 PTMScan antibodies, the flowthrough applied to the Rme2a PTMScan antibodies, and that flowthrough applied to Rme2s antibodies. Antibody epitopes are shown. Peptides were eluted and subject to mass spectrometry. A sample of input peptides was reserved for total proteome analysis **c.** Example ETD spectrum of the C-terminal peptide from small nuclear ribonuclearprotein Sm D1. The peptide fragment from residue 93 to 118 and contains 9x dimethylarginines, all site localized. The region from 350 to 750 m/z of the full mass spectrum (Left/Top) is expanded in the (right/bottom) spectra.

In our studies, we performed triplicate biological replicates in which we digested total proteome with trypsin and isolated total proteome tryptic peptides (**Supplemental Figures S3a,b**). We then performed sequential peptide immunoprecipitations with the Rme1 antibodies, followed by applying the flow-through to the Rme2a antibodies, and finally by applying the flow-through to the Rme2s antibodies (**Figure 3b**). Illustrating the biased specificity of the antibodies, the CST antigens for each antibody pool are shown. As described below, both the triplicate input peptides–representing the total proteome in each condition–as well as the immunoprecipitated proteins were subjected to mass spectrometry analysis. Parallel studies using the GluC enzyme resulted in spectra too complex to comprehensively analyze (*data not shown*).

The combination of the multiply modified peptides along with the trypsin digestion, sequential IPs, and cell treatment creates a complicated network of peptides. Therefore, we developed a decision-tree based mass spectrometry analysis that relied on ETD for the primary means of fragmentation for modified peptides. The Thermo Fusion Tribrid allows for sensitive and fast peptide analysis by utilizing a customized instrument method to make methylated peptides a priority. By allowing for charge-state based MS/MS analysis, shorter peptides were analyzed with the linear ion trap while more complex, longer peptides were analyzed with the Orbitrap. Additionally, ETD based fragmentation was optimal for confidently site-localizing modifications on the arginine-rich, tryptic peptides.

In our new approach, mass spectra were searched with a decision tree-based analysis utilizing a combination of Byonic by Protein Metrics and Proteome Discoverer 2.2. Proteome Discoverer filtered MS/MS by mass analyzer for data-base searching. Byonic enabled the database search to contain sufficient missed cleavages resulting from the modified arginine residues as well as adequate numbers of modifications.

The average total number of peptides identified in each PTMScan IP are shown in **Supplemental Figure S4a** (thousands for each IP). Demonstrating the specificity of the procedure, across all the IPs, approximately 75% of the total peptides contained methylation (**Supplemental Figure S4b**). As expected from the sequential immunoprecipitations, there were about 50% more Rme1-enriched peptides than Rme2s or Rme2a (**Supplemental Figure S4c**). The peptide abundances revealed that most peptides in each IP had either monomethylarginine or dimethylarginine. A smaller fraction of the peptide abundances exhibited a hybrid state, with both a monomethyl- and dimethylarginine (**Supplemental Figure S4d**).

To analyze the precipitated peptides and annotate methylated peptides, we plotted the log2 of peptide level drug-treatment abundance changes versus the log2 of the p-value (**Figure 4a-c**, Rme1, Rme2s, Rme2a IPs, respectively). As expected from the western blots, there were fewer overall peptide level abundance changes with GSK591 treatment (left green plots), while MS023-treatment (purple) resulted in dramatic increases in both Rme1- and Rme2s-containing peptides. Specific methylation-site level changes are shown in **Supplemental Sheet 1**. Based on the peptide data, we then assigned protein-level enrichment for each IP in each condition, showing 382 total proteins found in these IPs (**Supplemental Sheet 2**). For the protein enrichment in each condition, we calculated row z-scores and performed k-means clustering. As shown in **Figure 4d**, three major clusters were apparent. Consistent with known PRMT substrate scavenging, within each cluster the influence of each drug to alter the methylated proteins is clear. To understand the classes of proteins that are mono- or dimethylated, we performed ontology analysis and displayed the results using the semantic space similarity calculator REViGO (Supek et al. 2011) (**Figure 4e**). Consistent with prior studies, the methylated-protein ontologies were primarily related to nuclear function, transcription, splicing and translation. However, each enriched methylated state did show distinct GO enrichments. This suggests that unique molecular consequences for each methylation state are likely.

**Figure 4.**
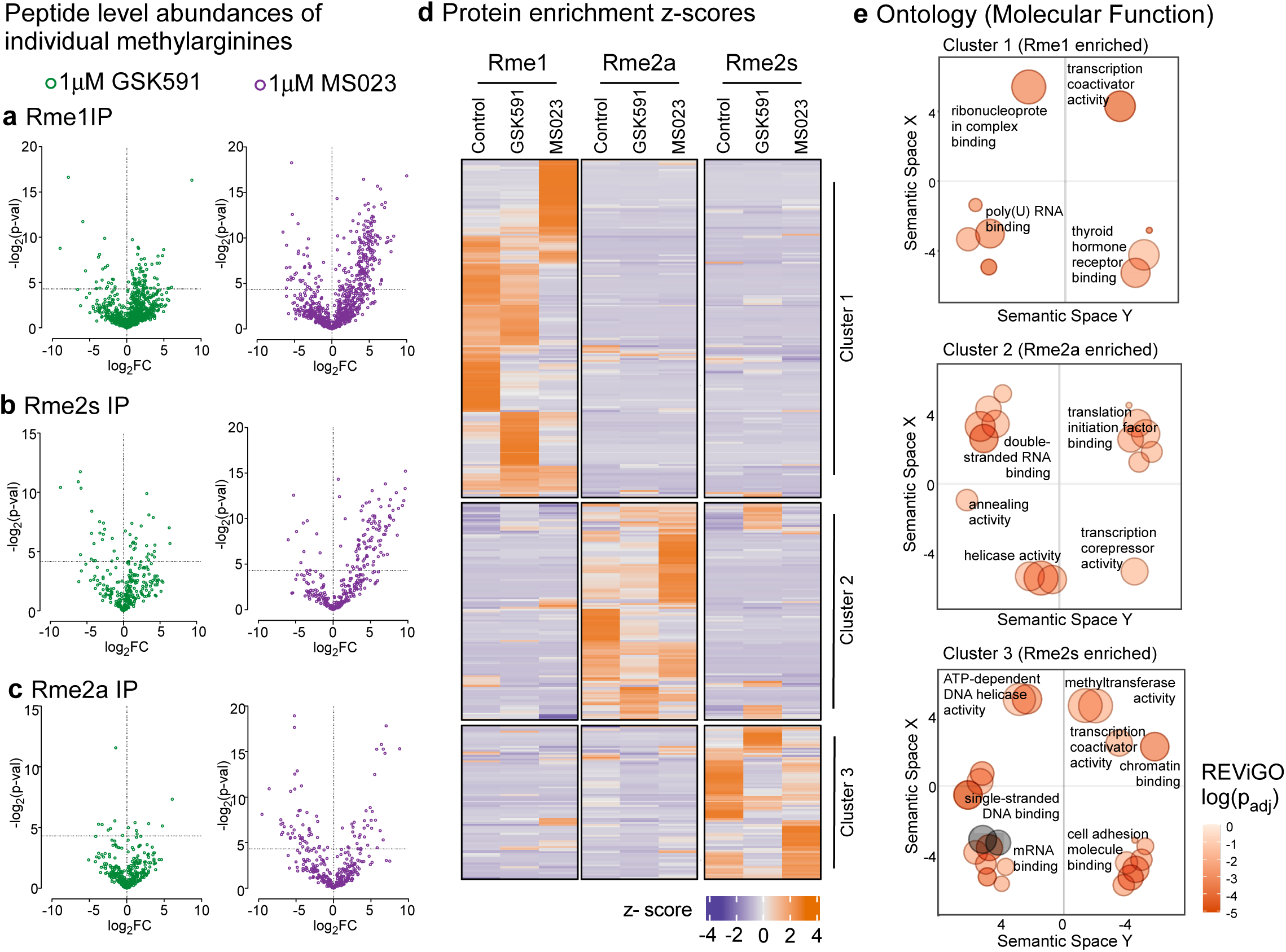
PTMScan site and protein analysis reveals PRMT-inhibitor dependent changes in arginine methylation. **a.** Volcano plot of monomethylarginine (Rme1) peptide enrichments for GSK591 (left, green) and MS023 (right, purple) treated cells. Log2 fold-change is on the x-axis, while the negative log2 of the p-value is shown on the y-axis (dashed x-axis line represents p ∼ .05). **b.** Volcano plot of symmetric dimethylarginine (Rme2s) peptide enrichments for GSK591 (left, green) and MS023 (right, purple) treated cells. **c.** Volcano plot of asymmetric dimethylarginine (Rme2a) peptide enrichments for GSK591 (left, green) and MS023 (right, purple) treated cells. **d.** 382 proteins found in all three IPs in all three conditions were clustered by row z-score (negative z-score shown in purple, positive z-score shown in orange). **e.** Each of the three protein clusters of proteins were analyzed for GO term enrichment. Shown is the semantic space REViGO plot for each of the molecular function groupings.

To further compare enriched methylarginine sites in the drug treatments, we plotted the log2 fold-change of the methylated arginine specific sites that were significantly altered in both GSK591 and MS023. As shown in **Supplemental Figure S5a**, almost all sites found in both datasets had positive correlations, either up- or down-regulated. This surprising observation is consistent with known substrate scavenging and could be due to the distributive nature of PRMT activities: any enzyme may catalyze Rme1 followed by symmetric or asymmetric dimethylation. Furthermore, the Rme1, Rme2s, and Rme2a enriched peptide motifs were almost entirely identical (**Supplemental Figure S5b**), consistent with the likely extreme bias of the antibody enrichment.

To better understand the types of proteins that are methylated in our PTMScan analysis, we first characterized the entire human proteome in terms of molecular weight, hydrophobicity, isoelectric point, and predicted intrinsic disorder. Using the RAPID algorithm (Yan et al. 2013), we probed 56,392 human Uniprot sequences, covering most known human protein variants. For each sequence, we also calculated the molecular weight, the predicted hydrophobicity as determined by GRAVY, and the predicted isoelectric point as calculated at isoelectricdb.org (Kozlowski 2016). We plotted the percent disorder, as calculated by RAPID, versus these values to illustrate the characteristics of the human proteome (**Figure 5a-c**). The median predicted disorder was 18.1%, the median molecular weight was 31.4 kDa, the median hydrophobicity was -0.37, and the median isoelectric point was 7.03. These plots gave us a unique perspective on the proteome: proteins of greater predicted intrinsic disorder are smaller than the median, substantially less hydrophobic, and more highly charged.

**Figure 5.**
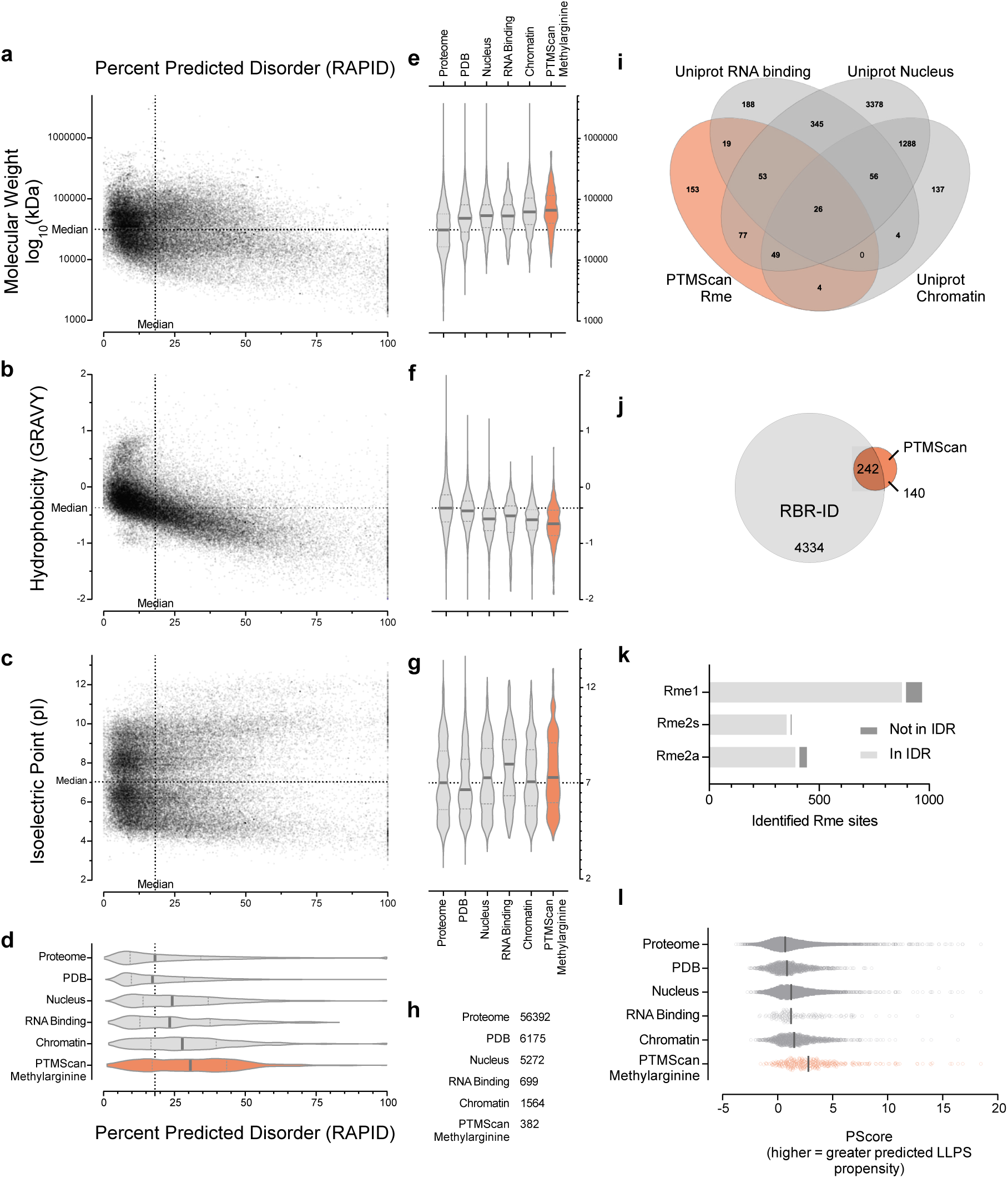
Proteome characteristics reveal the nature of the PTMscan arginine methylome. **a.** 56392 human proteins (Uniprot 2012) were plotted with their RAPID predicted intrinsic disorder percentage of the protein on the x-axis and the log10 of their molecular weight (Da) on the y-axis. The median intrinsic disorder across the proteome (18.1%) is indicated, as is the median molecular weight (31.4 kDa). **b.** The proteins were plotted with their RAPID predicted intrinsic disorder on the x-axis and their hydrophobicity as calculated by GRAVY on the y-axis. Positive scores are hydrophobic, while negative scores are hydrophilic. The median hydrophobicity across the proteome was -0.37. **c.** The proteins were plotted with their RAPID predicted intrinsic disorder on the x-axis and their isoelectric point on the y-axis. Positive scores are hydrophobic, while negative scores are hydrophilic. The median isoelectric point across the proteome was 7.03. **d.** The RAPID percent disorder distribution of the proteome, PDB, Nucleus, RNA-binding, chromatin, and methylarginine (orange) sets are shown as violin plots. The median is indicated with a dark line and the quartiles are shown with a dashed line. **e.** The molecular weight distribution of the proteome, PDB, Nucleus, RNA-binding, chromatin, and methylarginine (orange) sets are shown as violin plots as in d. **f.** The GRAVY hydrophobicity distribution of the proteome, PDB, Nucleus, RNA-binding, chromatin, and methylarginine (orange) sets are shown as violin plots as in d. **g.** The isoelectric point distribution of the proteome, PDB, Nucleus, RNA-binding, chromatin, and methylarginine (orange) sets are shown as violin plots as in d. **h.** Table showing the number of proteins in each set **i.** Venn diagram of the intersection between the PTMScan methylarginine containing proteins, RNA binding, Nucleus, and chromatin sets. 153 of the methylarginine containing proteins were not found in these sets. **j.** Venn diagram showing the intersection human proteins between the PTMScan methylarginine containing proteins and those previously identified to be bound to RNA using RBR-ID. 140 of the methylarginine containing proteins were not found in this set. **k.** Percent of methylated arginine residues found in intrinsically disordered regions (IDR, light gray) shown as a histogram: Rme1 (91%) in IDRs; Rme2s (95%); Rme2a (88%) **l.** The PScore distribution, indicating pi-pi mediated liquid-liquid phase separation (LLPS) propensity, is shown for the proteome, PDB, Nucleus, RNA-binding, chromatin, and methylarginine (orange) sets. The median is shown with a dark line. Across the figure, all distributions shown were significant (Kruskal-Wallis test, adjusted p-val < 0.01) as compared to the Uniprot total proteome, except for the pI of the chromatin fraction vs the proteome (n.s.)

For comparison with all 382 identified proteins containing methylarginine, we used Uniprot classifications to categorize human proteins, including proteins with solved PDB structures, those annotated to be found in the nucleus, those known to be involved in RNA binding, and proteins found associated with chromatin. In **Figure 5d-h**, we compared these distributions. The methylarginine proteins had the largest degree of predicted disorder but were substantially larger in MW and less hydrophobic than most. Importantly, methylarginine containing proteins had a wide charge distribution, consistent with enrichment of basic patches (e.g. “GR”) and neighboring acidic stretches. We intersected these protein groups in a Venn diagram (**Figure 5i**); this illustrated that while the majority of the identified methylarginine proteins fell in the nucleus, chromatin, or RNA-binding categories, 153 of them did not. To further confirm the likelihood that these methylated proteins are involved in RNA binding, we intersected the proteins with a set of human proteins positively identified through RBR-ID as interacting with RNA (He et al. 2016). As shown in **Figure 5j**, 242 of the proteins intersected. These observations are all consistent.

As methylarginine has been hypothesized to potentially be involved in regulated liquid-liquid phase separation (LLPS), we further tested properties of the methylated proteins. First, as many LLPS and nucleic acid interactions are mediated through intrinsically disordered regions (IDRs), we asked if the identified methylarginine sites were embedded in IDRs. We probed the MobiDB database (Piovesan et al. 2017) (https://mobidb.bio.unipd.it/) of IDRs and interrogated if each site was in an annotated IDR. As shown in **Figure 5k**, roughly 95% of the identified sites in all three IPs were found in IDRs. Finally, we used the PScore algorithm to predict LLPS propensity; this algorithm specifically probes putative pi-pi contacts, the kind that may be directly influenced by arginine methylation (Vernon et al. 2018). As shown in **Figure 5l**, the methylarginine containing proteins had a significantly higher PScore and therefore a higher predicted propensity for LLPS.

### Biological consequences of PRMT inhibition

To begin to understand the functional consequences of PRMT activity and of the inhibition of their activity, we probed both the proteome and the transcriptome of GSK591- and MS023-treated A549 cells. Note that extensive analysis of these consequences will be described elsewhere (*Maron, Burgos, Shechter, et al, in preparation*, and *Chen, Shechter, et al, in preparation*). PRMTs have previously been shown to alter both transcription and translation, so we wanted to determine if these consequences were related or were independent.

We first used the input proteome tryptic peptides from the PTMScan–triplicate in each of DMSO, GSK591, and MS023 treatments–and calculated proteome up- and down changes. Relative changes in protein abundance was estimated after performing log2 transformation and normalization by the average quantitative value for each sample. This transformation avoided biases in total protein injection amount, hence the symmetric volcano plots. For both conditions, the log2-fold change of the drug-treatment compared to the DMSO control is plotted against the log2 of the p-value in **Figure 6a,b**. The p-value was obtained by performing a heteroscedastic t-test between drug treatment and DMSO control. Out of 4141 proteins confidently observed in all conditions, we observed 267 significantly (p<0.05) altered proteins in the GSK591-treated samples and 324 significantly altered proteins in the MS023-treated samples (**Supplemental Sheet 3)**.

**Figure 6.**
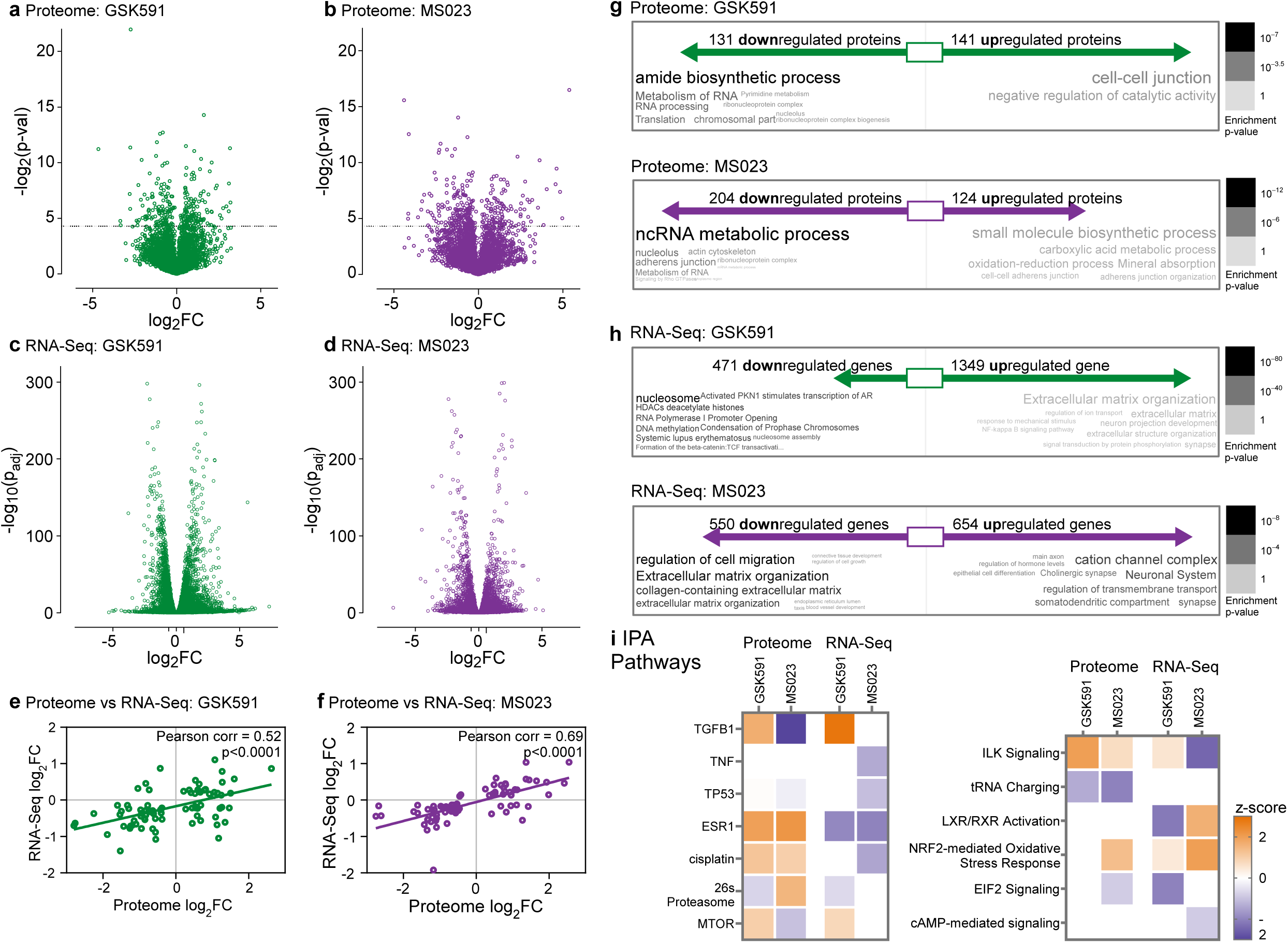
Proteomic and RNA-Seq analysis reveal overlapping and independent cellular regulation by PRMT inhibitors. **a.** Volcano plot of the GSK591-regulated total proteome. Log2 of the protein fold-change is shown on the x-axis and negative log2 of the p-value is shown on the y-axis **b.** Volcano plot of the MS023-regulated total proteome. Log2 of the protein fold-change is shown on the x-axis and negative log2 of the p-value is shown on the y-axis **c.** Volcano plot of the GSK591-regulated transcriptome. Log2 of the protein fold-change is shown on the x-axis and negative log10 of the adjusted p-value is shown on the y-axis **d.** Volcano plot of the MS023-regulated transcriptome. Log2 of the protein fold-change is shown on the x-axis and negative log10 of the adjusted p-value is shown on the y-axis **e.** Correlation of GSK591 significantly altered transcripts and their corresponding proteins are shown. Pearson correlation is inset. **f.** Correlation of MS023 significantly altered transcripts and their corresponding proteins are shown. Pearson correlation is inset. **g.** GOsummaries word clouds for significantly up and down regulated proteins (p<0.05) are shown in green for GSK591 (top panel) and in purple for MS023 (bottom panel). The enrichment p-value is indicated by the words’ gray coloring. **h.** GOsummaries word clouds for significantly up and down regulated transcripts (log2FC>|1| and padj < 0.01) are shown in green for GSK591 (top panel) and in purple for MS023 (bottom panel). The enrichment p-value is indicated by the words’ gray coloring. **i.** Selected IPA analysis upstream pathway (left panel) and canonical pathway (right panel) are shown as z-score heatmaps (positive z-score shown in orange, negative z-score shown in blue).

Next, we performed RNA sequencing on A549 cells similarly treated for 1 week with DMSO, GSK591, or MS023. As shown in **Figure 6c,d**, thousands of transcripts were up- and down-regulated in both PRMT-drug treatments. As noted above, more specific transcriptome consequences will be described elsewhere.

Through histone and non-histone methylation, PRMTs may independently regulate transcription and translation. To test if there was concordance between altered protein expression and altered transcription, we identified the 79 genes and corresponding proteins that were significantly altered in both the proteome and RNA-Seq GSK591-datasets and 76 genes and corresponding proteins significantly altered in both MS023-datasets. As shown in **Figure 6e,f**, there was some correlation between the RNA-Seq and the proteome, particularly for the MS023 treatment (Pearson correlation = 0.69). However, while GSK591 treatment still had a general positive correlation (Pearson correlation = 0.52), there were more transcripts and proteins that had opposite transcript and protein regulation.

To identify the cellular and molecular consequences of these transcriptional and proteomic regulatory events, we identified <u>g</u>ene <u>o</u>ntology (GO) term enrichments and, using the GOSummaries R package, we plotted them as differential word clouds (**Figure 6g,h**). As expected from the general correlation of proteome and transcriptome, the up- and down-regulated GO enrichments were generally similar (**Supplemental Figure S6**). For both, these included terms regarding nuclear and RNA function as well as terms regarding extracellular matrix and cellular adhesion. However, specific differences were apparent and are further illustrated using IPA pathway analysis (**Figure 6i**). As a positive control consistent with our previous studies on regulation of the TGFβ pathway by PRMT5 (Chen et al. 2017), GSK591 caused an upregulation of TGFβ1 upstream pathway in both the proteome and transcriptome. In contrast, the ESR1 estrogen receptor pathway exhibited proteome upregulation in both drugs, but transcriptome downregulation in both conditions.

As we collected three types of proteomic and transcriptomic data of PRMT-regulated events, we aimed to compare their consequences. To transform the analysis to comparable data, we used the GOSemSim R package to both identify enriched GO terms and build a similarity matrix. As shown in **Figure 7a** and **Supplemental Figure S7**, we probed the similarity of ontologies for GSK591- and MS023-up and down regulated transcripts and proteins, as well as for the Rme1, Rme2s, and Rme2a enriched proteins. While the GO terms were generally similar between all three classes of data–and interestingly the RNA-Seq was most similar to itself in all conditions–differences were apparent. Particularly in the “Biological Pathway” and “Molecular Function” umbrella terms, the proteome and PTM-Scan had more concordance than they did with the transcriptome. The monomethylarginine IP dataset, in particular, was more similar in ontology to the proteome than to the other IP sets. This suggests a possible connection between monomethylarginine and protein turnover and/or translation (**Figure 7b**).

**Figure 7.**
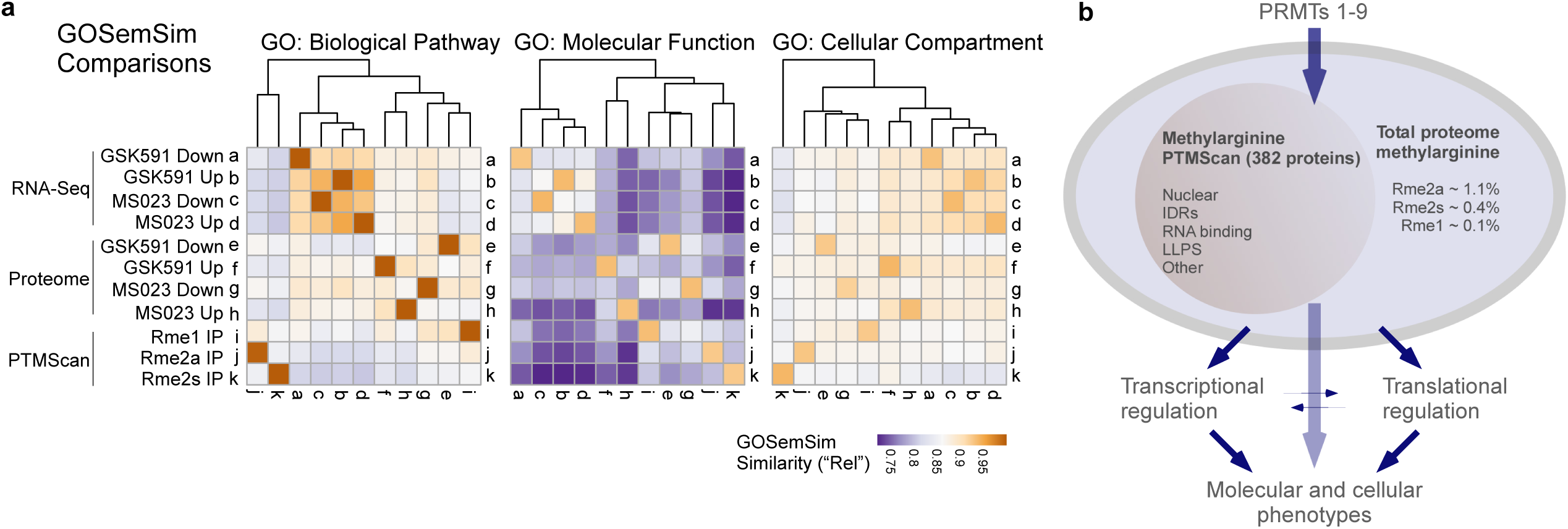
Summary of the results reveals that PTMScan only captures a subset of the cellular regulation by the family of PRMTs. **a.** GOSemSim matrices of gene ontology comparisons between the RNA-Seq, proteomic, and PTMScan altered genes and proteins. Biological pathway, Molecular Function, and Cellular Compartment ontology similarities, using the “Rel” semantic comparison metric, are shown as a heatmap. **b.** Cartoon representation of our findings, exhibiting that PRMTs methylate up to 4% of the total proteome arginine and that our PTMscan methylarginine analysis only captured a subset of these proteins, enriched in IDRs, nuclear function, RNA binding, and potentially LLPS. We did show that PRMTs regulate both transcription and translation.

## Discussion

In this study, our goal was to provide a comprehensive understanding of the consequences of PRMT activity and cellular methylarginine on the transcriptome and proteome of human cells. As highly-effective and specific chemical probes have been developed for the main classes of PRMT enzymes, to test cellular consequences of the loss of these enzymes we used MS023 (a general inhibitor of most Type I enzymes)(Eram et al. 2016) and GSK591 (a specific PRMT5 inhibitor, also known as EPZ015866 or GSK3203591)(Duncan et al. 2015). We developed both new robust mass spectrometry methods for characterizing the total abundance of methylarginine species and a label-free methodology for PTMScan peptide immunoaffinity analysis. In parallel, to determine PRMT substrate specificity, we utilized a new *in vitro* methyltransferase assay against degenerate peptides. Finally, to understand how PRMT activities against these cellular substrates regulates the transcriptome and proteome, we performed RNA-Seq and total proteome analysis in control and drug-treated conditions. Altogether, our work is the first to comprehensively integrate understanding of PRMT substrates with the transcriptomic and proteomic consequences of their activity.

Multiple approaches have been previously used to characterize total proteome arginine methylation; most relied upon hydrolyzed-protein reversed-phase chromatography coupled with standards and peak integration. Here, after total protein acid hydrolysis to produce constituent amino acids, we developed a new procedure. To more precisely characterize the specific methylarginine isoforms, we used high resolution UHPLC chromatography and mass spectrometry. Fragment ion losses were characterized for each methylarginine species. Calibration curves with ^1^H-NMR-calibrated amino acid standards allowed us to accurately determine the biological methylarginine abundances. Using this new approach, we showed that methylarginine is more abundant than previously shown. Importantly, we also showed that GSK591-dependent loss of PRMT5 activity led to a substantial reduction in Rme2s, but MS023 treatment only lead to a 50% reduction in Rme2a. Since we used both drugs at relatively high concentrations (10-100X) compared to their IC50s, there are two possible explanations for these observations. Methylarginine half-life is likely long, suggesting that even after one week of treatment, some residual long-lived methylation may be present. Future studies will need to use SILAC type approaches to determine methylarginine half-lives (Zee et al. 2010). Alternatively, other PRMTs with lower sensitivity to MS023–such as PRMT2 and PRMT3 and the untargeted in this study Type III enzyme PRMT7–may produce these methylations. Importantly, the large and poorly documented family of methyltransferase-like (METTL) SAM-dependent methyltransferases includes METTL23, recently demonstrated to catalyze histone arginine methylation (Hatanaka et al. 2017). Consistent with our observations, this suggests that other non-PRMT enzymes could catalyze arginine methylation. Future studies are necessary to test this possibility.

Our studies also included an advance of the previously performed OPAL (oriented peptide array library) technology. In this work, we used more quantitative flashplate scintillation counting of PRMT activity towards immobilized degenerate peptides (Cornett et al. 2018). As these peptides contained a fixed central arginine substrate along with a single other fixed amino acid position (excluding Cysteine), we were able to derive *in vitro* substrate preferences for the three most abundant enzymes: PRMT1, PRMT4/CARM1, and PRMT5. Our results showed a wider substrate preference for PRMT1 and PRMT5 than previously known, including some likely modulatory consequences of the MEP50 substrate presenter (Ho et al. 2013; Burgos et al. 2015). Most strikingly, CARM1 showed a remarkably distinct substrate consensus, with marked preference for hydrophobic substrates. While this was previously hinted at (Gayatri et al. 2016; Shishkova et al. 2017), our evidence should prompt a deeper search for unique CARM1 substrates in cells.

To characterize the cellular PRMT substrates, we employed the widely used PTMScan approach. This approach relies on proteolysis of the proteome to ensure soluble peptides for targeted immunoprecipitation. Uniquely, we performed this experiment in three conditions with three targeted antigens: in control, GSK591-, or MS023-treated cells with antibodies against Rme1, Rme2a, and Rme2s. To make the experiment time- and cost-effective, we performed successive immunoprecipitations. We developed a new method for the detection of methylarginine containing peptides by LC-MS/MS that relied on a decision tree-based instrument method and database search. The method, which prioritized ETD fragmentation over collision-based fragmentation, allowed for confident site localization of mono- and di- methylated residues in multiply modified peptides. With this approach, peptides with up to 9 modified arginine residues were able to be identified. The decision tree-based instrument method enabled the best fragmentation method to be applied for each peptide. ETD, a charge dependent ion/ion reaction, was used for arginine rich peptides; collision-based HCD was implemented for peptides with fewer missed cleavages. Additionally, the decision tree-based method took advantage of both mass analyzers on the Fusion Tribrid mass spectrometer, with the high resolution Orbitrap reserved for the most complex (arginine-rich) peptides. Furthermore, the database search allowed for up to 12 missed cleavages. As previous methods have been limited to collision-based fragmentation and were only searched with 3 or 4 missed cleavages (Larsen et al. 2016; Musiani et al. 2019), our approach is a substantial advance.

However, the PTMScan approach is still limited by both the protease used for digestion and by the bias of the employed antibodies. While we attempted to overcome the first limitation by employing both trypsin and GluC digestion, the GluC-peptides were much larger with less consistent enzymatic cleavage and with many modifications were too complex for quantitative study. Therefore, we only used the tryptic peptides. As with others who have used this approach, the experimental design still has the potential to create a complicated web of results. The results returned from search algorithms include summed abundances at the protein level and at the peptide level. As the search algorithm used intact mass to distinguish peptides, the same peptide may be returned with varying numbers of methylations. Thus, the same peptide may look depleted in a treated sample compared to the control only to see the same peptide with fewer methyl groups increased compared to the control. At the protein level, the search algorithm will return results summed for the protein. While a result may be reported as the same for a treated sample, it could be that a region of methylated peptides are missing compared to the control, but this would not be reflected on the summed protein level.

Our study did reveal important insights: MS023 treatment, inhibiting the Type I enzymes, increased the abundance of recognized Rme1 substrates, implying a putative role for PRMT5 in catalyzing much of the first methylation step; consistent with the total proteome analysis conducted in Figure 1, most of the Rme2s substrate signals were lost upon GSK591 treatment; and, consistent with known PRMT substrate scavenging, both drug treatments caused changes in Rme2a substrates. However, each methylated state did show distinct GO enrichments. This suggests that unique molecular consequences for each methylation state are likely.

Consistent with similar work, many of our enriched methylated proteins were found in RNA-binding and nuclear function. As we had prior interest in protein intrinsic disorder (Warren and Shechter 2017), we hypothesized that many methylarginine-containing proteins were embedded in IDRs. We produced a unique analysis of the total human proteome, showing unique characteristics of proteins containing predicted intrinsic disorder. Consistently, the proteins we identified as containing methylarginine were significantly enriched in intrinsic disorder. However, unlike most intrinsically disordered proteins, methylarginine-containing proteins were larger than the median and less hydrophobic. These characteristics suggest a potential unique role in charged- and disorder based regulatory function. We demonstrated that the methylarginine-containing proteins were also more likely to be found in the nucleus and directly bind RNA; these features are commonly enriched in proteins driving liquid-liquid phase separation (LLPS) and molecular condensates (Ditlev et al. 2018). Consistent with a model of methylarginine potentially regulating LLPS, the PScore algorithm we demonstrated that methylarginine-containing proteins were significantly more likely drive this process. Given the total picture of methylarginine enrichment in nuclear- and RNA function, and the abundant new literature on how LLPS is critical for nuclear function (Strom and Brangwynne 2019), it is compelling to hypothesize that specific methylarginines directly regulate these events. Future studies will directly test this hypothesis.

In line with the evidence above that protein arginine methylation is enriched-in and may regulate nuclear and nucleic acid biology, it was quite striking that the transcriptome and proteome regulated by the inhibition of both Type I- and Type II-PRMTs was characterized primarily by downstream cellular consequences. These included GO terms regarding extracellular matrix, metabolism, and biosynthetic processes. As the methylation of proteins may have multiple consequences, including direct regulation of transcription and regulation of translation, this suggests a nested PRMT regulatory regime. Consistent with this model, we were able to compare the ontologies of all three independent datasets: the RNA-Seq of PRMT inhibitor up- and down-regulated genes; the proteome in the same conditions; and the Rme1, Rme2a, and Rme2s PTMscan proteins. Clustering of these datasets showed that, generally, the transcriptome gene ontologies grouped together while the proteome and the methylarginine ontologies clustered together. A simple explanation for this observation is that the transcriptome is independently regulated–perhaps by histone arginine methylation– while subsequent translation is more closely regulated with actual protein function.

Altogether, our new approaches of total amino acid analysis by mass spectrometry, in vitro PRMT activity assays, robust label-free PTMScan, proteomics, and transcriptomics demonstrate that protein arginine methylation is more complex and more abundant than previously known. Future work will be necessary to assign specific functions to all three methylarginine states, to identify more enzymes responsible for arginine methylation, and to understand the cellular role of PRMTs.

## Experimental methods

### Cell Culture and treatment with PRMTs Inhibitors

A549 cells (freshly purchased for this study from ATCC) were cultured in DMEM medium (FBS and Penicillin/Streptomycin at 10% and 1X final concentration, respectively), either exposed to MS023 Type-I PRMT inhibitor or GSK591 PRMT5 specific inhibitor (1 µM final concentration) for seven days; both compounds were obtained from Cayman Chemicals. 150 mm plates (30 mL medium) were seeded with 0.2 x 10^5^ and 0.4 x 10^5^ cells for the control and drug-treated groups, respectively. Incubator maintained temperature and CO_2_ concentration at 37 °C and 5%, respectively. Following trypsin digestion, cells were pelleted via centrifugation (300g, 3 min) and further washed with PBS buffer.

### Isolation of Whole Protein Samples and Acid-Hydrolysis

Per group, a total of 10^7^ cells were isolated and flash frozen. On ice, A549 cells pellets (50 mL tube) were resuspended in 4 mL freshly prepared Lysis Buffer (150 mM NaHCO_3_, 8 M Urea, supplemented with 1 mM DTT; pH 8.3) to obtain a clear lysate. Trichloroacetic acid solution–made from 10g TCA dissolved into 4.5 mL of water; i.e. 100% TCA solution or 100 g per 100 mL solution−was added to lysate (1:4 v/v) to precipitate proteins. Whole protein pellet was isolated by centrifugation, further washed with ice-cold acetone (5X) and dry under vacuum. Dry pellets (∼25−50 mg) were transferred into glass pressure tube (AceGlass #8648-230), flushed with argon prior to adding hydrochloric acid (3 mL of 6 M sequencing grade solution; Thermo Scientific #PI24308). Sealed tubes were kept at 125° C for 48h (oil bath) to achieve acid-hydrolysis of the whole A549 proteome. Light yellow solutions were further diluted (roughly 5000-fold) to fit in the calibration curve prior to MS analysis/quantitation.

### Quantitation of Arginine and Lysine Species within Acid-hydrolyzed A549 Whole Proteome

Amino acid mixture was resolved onto an Agilent Zorbax Eclipse Plus C18 RRHD column (2.1×150 mm, 1.8 µm; kept at 40C) using the mobile phases A 0.1% formic acid (FA) in water and B 0.1% FA in acetonitrile, following an optimized gradient. Within the Agilent 1290 UPLC system, flow rate (400 µL min^-1^) and injection volume (10 µL; auto-sampler at 10C) were kept constant. Arginine species standards (i.e. R, Rme1, Rme2s, Rme2a; concentration determined by ^1^H NMR using adenosine internal standard, δ 6.1 ppm, d, ^1^H) were identified and quantitated using the Agilent 6490 triple quadrupole (QqQ) mass spectrometer system under a positive electrospray ionization mode. While a Low Resolution analysis was performed, several benchmarks were performed to ensure sensitivity and accuracy of our study: 1) the liquid chromatography gradient was optimized to maximize compound’s separation and peak width for sensitivity, 2) all standards’ molecular weight were confirmed by MS (no substantial adducts present; e.g. sodium), 3) MS/MS were performed to confirm identity of each compound and to identify unique product ions for MRM transitions, 4) collision energies were optimized for each compounds MRM transition (at least 2 per compound; at times sensitivity was sacrificed for specificity), 5) dynamic range and sensitivity limits for each MRM transition (mainly quantifier transition) were established, 6) when needed, the amount of contribution of one transition to another was established to prevent decreased specificity and sensitivity and 7) calibration curves were repeated in between sample measurements. Set-up: source nitrogen gas temperature set at 290C with a flow rate of 16 L min-1; nebulizer 45 psi; sheath gas temperature set at 250C with a flow rate of 11 L min-1; capillary at 4,000 V. The MRM transitions each with a 50 msec dwell time were (collision energy in parentheses with **Q** identifying the **quantifier**): R, 175.1 (175.1190) to **70** (**70.0652**; 3,4-dihydro-2H-pyrrol-1-ium) m/z (25 V; Q), 175.1 (175.1190) to 60 (60.0557) m/z (10 V); Rme1, 189.1 (189.1347) to **70** (**70.0652**) m/z (25 V; Q), 189.1 (189.1347) to 74 (74.0713) m/z (10 V); Rme2s, 203.2 (203.1503) to **172** (**172.1081**) m/z (5 V; Q), 203.2 (203.1503) to 133 (133.0972) m/z (5 V); Rme2a, 203.2 (203.1503) to **46** (**46.0652**) m/z (10 V; Q), 203.2 (203.1503) to 112 (112.0870) m/z (10 V).

### Oriented Peptide Array Library (OPAL) and PRMT screening

OPAL peptides were synthesized in the following fashion: A-ZXX-R-XXX-A-PEG-Biotin, A-XZX-R-XXX-A-PEG-Biotin, A-XXZ-R-XXX-A-PEG-Biotin, A-XXX-R-ZXX-A-PEG-Biotin, A-XXX-R-XZX-A-PEG-Biotin, A-ZXX-R-XXZ-A-PEG-Biotin in which R is a fixed Arginine, X is any amino acid except cysteine, and Z is a fixed position. FlashPlate wells were coated with Streptavidin. Biotinylated peptide pools were treated with recombinant *Hs*PRMT1, *Hs*PRMT4/CARM1, *Ce*PRMT5, or *Hs*PRMT5-MEP50 (1 µg of peptide, 50mM Tris-HCl pH 8.8, 5 mM MgCl_2_, 4 mM DTT, 1 µCi ^3^H-SAM and ∼1 µg of enzyme) for 1 hour at 30° C following which they are transferred to FlashPlates (Perkin Elmer) and incubated for 30 min for peptide capture. Polystyrenebased scintillant was added. The radioactivity of the biotinylated peptides, now bound to the wells via streptavidin, was counted. Unbound SAM which is still in the solution in the well is not counted. To generate the heatmap, we collected MicroBeta counts per minute (CPM). For each position (P3- to P3+), all the readings were normalized to the highest value so that the highest CPM reading has a value of 1 and all others are expressed as a fraction of the highest CPM. The assay was performed in triplicate.

### PTMScan® peptides and immunoprecipitation

PTMScan was conducted entirely according to the manufacturer’s instructions (Cell Signaling Technology, #13563). Briefly, 2 x 10^8^ A549 cells of each treated condition were washed with PBS and scraped freshly made room temperature in urea lysis buffer (20 mM HEPES, pH 8.0, 9 M Urea, 1 mM sodium orthovanadate, 2.5 mM sodium pyrophosphate, 1 mM β-glycerophosphate). Cell lysates were sonicated and centrifuged. The diluted soluble protein lysate (to a final concentration of 2M Urea, 20mM HEPES pH 8.0) was reduced with DTT and alkylated with iodoacetimide. Each lysate was mixed digested overnight at room temperature with 100 µg Trypsin in 1mM HCl (Pierce 90305). GluC treatment was also performed and PTMScan conducted on these peptides; however, they were not used in the final analysis. Digestion was confirmed by SDS-PAGE. Digested peptides were acidified to 1% TFA and purified on SepPak C18 gravity columns (Waters WAT051910). Eluted peptides were lyophilized and dissolved in the manufacturer’s IAP buffer. A fraction of eluted peptides was reserved for the “Input proteome” analysis. Peptides were subjected to successive immunoprecipitations with the Rme1 (CST Kit # 12235), Rme2a (CST Kit # 13474), and Rme2s (CST Kit # 13563) prebound resins. Flowthrough from each IP was applied to the next resin. Resin was washed and eluted with two applications of 0.15% TFA.

### PTMScan mass spectrometric methods

Digested input proteome samples and and input PTMScan samples were desalted using C18 STAGE tips (3M). Proteome samples were acidified with 1% trifluoroacetic acid (TFA) prior to loading onto STAGE tip discs. The samples were washed with 0.1% TFA and eluted with 50% acetonitrile, 0.1% formic acid. Samples were dried and re-constituted with 0.1% formic acid.

Input samples were loaded into a Dionex UltiMate 3000 (Thermo Fisher Scientific) liquid chromatography system, coupled with an Q-Exactive HF instrument (Thermo Fisher Scientific). Peptides were separated on an in-house made capillary column (20 cm x 75 um fused silica column with a laser pulled tip, packed with Dr. Maisch, Reprosil-Pur 120 Å C18-AQ, 3 µm particles. The peptides were separated at 400 nL/min as follows: 3% solvent B (above) to 30% in 105 minutes, 50% solvent B at 120 minutes followed by column wash and re-equilibration.

The instrument method had a full MS scan with 120,000 resolution at 200 m/z and a 1,000,000 AGC target, acquired from 300 to 1500 m/z. The top 20 precursors from z = +2 to +6 were selected for isolation with a 2.0 m/z window for HCD fragmentation with NCE of 28. The MS/MS had a 200,000 AGC target and a 15,000 resolution at 200 m/z with a 30 second dynamic exclusion.

PTMScan samples were loaded into a Dionex UltiMate 3000 (Thermo Fisher Scientific) liquid chromatography system, in-line with an Orbitrap Fusion Tribrid mass spectrometer with ETD capabilities (Thermo Fisher Scientific). Peptides were separated on an in-house made capillary column (20 cm x 75 um fused silica column with a laser pulled tip, packed with Dr. Maisch, Reprosil-Pur 120 Å C18-AQ, 3 µm particles. Immunoprecipitated samples were analyzed with 300 nL/min flow rate as detailed: (solvent A: 0.1% formic acid; solvent B: 0.1% formic acid in 95% acetonitrile) 2% to 40% solvent B in 65 minutes, increased to 60% solvent B in 74 minutes, followed by column wash and re-equilibration.

The instrument method included a 60,000 resolution MS1 scan from 300 to 1500 m/z and allowed for a three second cycle time. Selection of precursors for MS/MS followed a decision tree approach based on charge state and precursor m/z range: 1.) Precursors with a z = +3 to +4, from 300 to 850 m/z were fragmented by ETD and analyzed in the ion trap. The AGC target was set to 20,000 with a maximum ion acquisition time of 100 ms. 2.) Precursors with a z = +5 to +12, from 300 to 850 m/z were fragmented by ETD and analyzed in the Orbitrap with 7,500 resolution. The AGC target was set to 100,000 with a maximum ion acquisition time of 200 ms. 3.) Precursors with a z = +2 to +4, from 300 to 1500 m/z were fragmented by HCD with a normalized collision energy (NCE) of 29 and analyzed in the ion trap. The AGC target was set to 20,000 with a maximum ion acquisition time of 75 ms. 4.) Precursors with a z = +5 to +8, from 850 to 1500 m/z were fragmented by HCD, with NCE of 29 and acquired in the Orbitrap with 7500 resolution at 200 m/z. The AGC target was set to 100,000 with a maximum ion acquisition time of 200 ms. A dynamic exclusion of 20 seconds was used for all precursors, as well as a 2.0 m/z isolation window.

### Database Search: PTMScan

The data was analyzed using Byonic (Protein Metrics) in Proteome Discoverer 2.3 (Thermo Fisher Scientific). Peptides identified through Proteome Discoverer were quantified using the area of their respective extracted ion chromatogram. Data were then log2-transformed and normalized by the average quantitative value of each sample to eliminate biases in sample recovery and pipetting. Before quantifying differences in relative abundances between samples, the intensity of the methylated peptide was normalized by the estimated abundance of its respective protein. This last step was performed to avoid changes in protein translation being confused with changes in methylation level. The PTMScan data was analyzed with the reviewed human proteome from SWISS-PROT, downloaded 17/09/2019 with 20,353 entries. Results were search allowing for up to 12 missed cleavages. The following modifications were allowed: carbamidomethyl on cysteine as fixed; dimethylation on arginine (common, up to 10) and lysine (rare, up to 2), methylation on arginine (up to 5) and lysine (common, up to 2), acetylation on protein n-termini and lysine (rare, up to 1), oxidation on methionine (rare, up to 2). The results were filtered for a 1% false discovery rate.

### Input Samples

The data was analyzed using Byonic (Protein Metrics) in Proteome Discoverer 2.3 (Thermo Fisher Scientific). The data was analyzed twice with separate two databases, the full reviewed human proteome from SWISS-PROT as per above as well as a database created from the RNA-Seq results. The search allowed for up to 4 missed cleavages and the following modifications: carbamidomethyl on cysteine as a fixed modification, acetylation on protein N-terminus, and up to two oxidized methionine residues per peptide. Identified spectra were filtered to a 1% false discovery rate using Percolator.

### Processing of database search result tables

The output of Proteome Discoverer (v2.3, Thermo Scientific) was processed entirely using Microsoft Excel. Briefly, total methylation enrichment was assessed by summing the intensity of all peptides carrying a methylation divided by the sum of all peptide intensities for each sample. Peptides were considered as “hybrid peptides” if they contained at least one monomethyl and one dimethyl group. For enrichment analysis, data were first log2 transformed, then normalized by subtracting the average of the values for each sample. The input proteome was subjected to the same normalization, and methylated peptides were then normalized by their respective protein abundance by subtracting the protein abundance from the peptide abundance. Statistical enrichment was assessed by using a two-tails heteroscedastic t-test (significant if p-value <0.05). For hierarchical clustering, data were first z-score normalized. For motif analysis, the position of the modification within each protein sequence was extrapolated, and we retrieved the sequence around the modified residue ±10 amino acids. To determine if methylarginine sites were embedded in intrinsically disordered domains,

### Proteome characteristic analysis

56392 human protein sequences (Uniprot 2012) and their predicted intrinsic disorder were obtained from http://biomine.cs.vcu.edu/servers/RAPID/homosapiens.txt (Yan et al. 2013) and molecular weight and isoelectric point were obtained from http://isoelectricpointdb.org/40/UP000005640_9606_all_isoelectric_point_proteome_Homo_sapiens_Human.html (Kozlowski 2016).

### RNA-Seq

For each condition, triplicate total RNA was extracted with mRNeasy kits (Qiagen), rRNA removed with Riboerase (Kapa Biosystems) and paired-end libraries were prepared with random hexamers (Kapa). Each library was sequenced to attain approximately 40M paired-end 150bp reads on a NextSeq 500. Sequences were mapped to hg19 using STAR (Dobin et al. 2012) and differential gene expression was determined using featureCounts and DESeq2 (Love et al. 2014).

### Bioinformatics and graphics

The following R packages were used in this study: ggSeqLogo was used to create the weblogo plots (Wagih 2017); clusterProfiler for determining gene ontology (Yu et al. 2012); and REViGO (Supek et al. 2011) and GOSemSim (Yu 2020) were used for ontology similarity analysis. All charts and histograms were plotted in Graphpad Prism v8, while final figures were assembled in Canvas X 2020.

## Supporting information

Lehman et al Supplemental Sheet 1

Lehman et al Supplemental Sheet 2

Lehman et al Supplemental Sheet 3

## Acknowledgments

This work was supported by the National Institutes of Health: R01GM108646 to D.S., R01GM037537 to D.F.H, R01GM126421 to M.T.B., and R44GM112234 to Z.W.S. D.S. was also supported by the American Lung Association Discovery Award LCD-564723. Thank you to Benjamin A. Garcia for sharing resources and instruments. Thank you to Protein Metrics for generously providing the Byonic search algorithm which was used in this work. We appreciate early access to PRMT inhibitors from the Structural Genomics Consortium (Toronto).

## Author contributions

S.M.L. developed mass spectrometry techniques and performed experiments; H.C. performed experiments; E.S.B. and E.N. developed and performed total arginine analysis; E.S.B. also expressed and purified *Ce*PRMT5 and *Xl*PRMT5-MEP50 enzymes; M.M. performed bioinformatic analysis; S.G. and M.T.B. developed and performed OPAL experiments; M.R.M, J.R.B, and Z.W.S. designed and produced the OPAL library; V.G. performed RNA-Seq analysis; D.L.B. and S.S. performed mass spectrometry computational analysis; J.S. and D.F.H. supervised mass spectrometry; D.S. supervised the study, performed bioinformatic analysis, and wrote the manuscript.

## Accession numbers

RNA-Seq data is deposited under GEO (*pending*). Mass spectrometry data is deposited under Chorus Project (1671).

## Supplemental Figure Legends

**Supplemental Figure S1.**
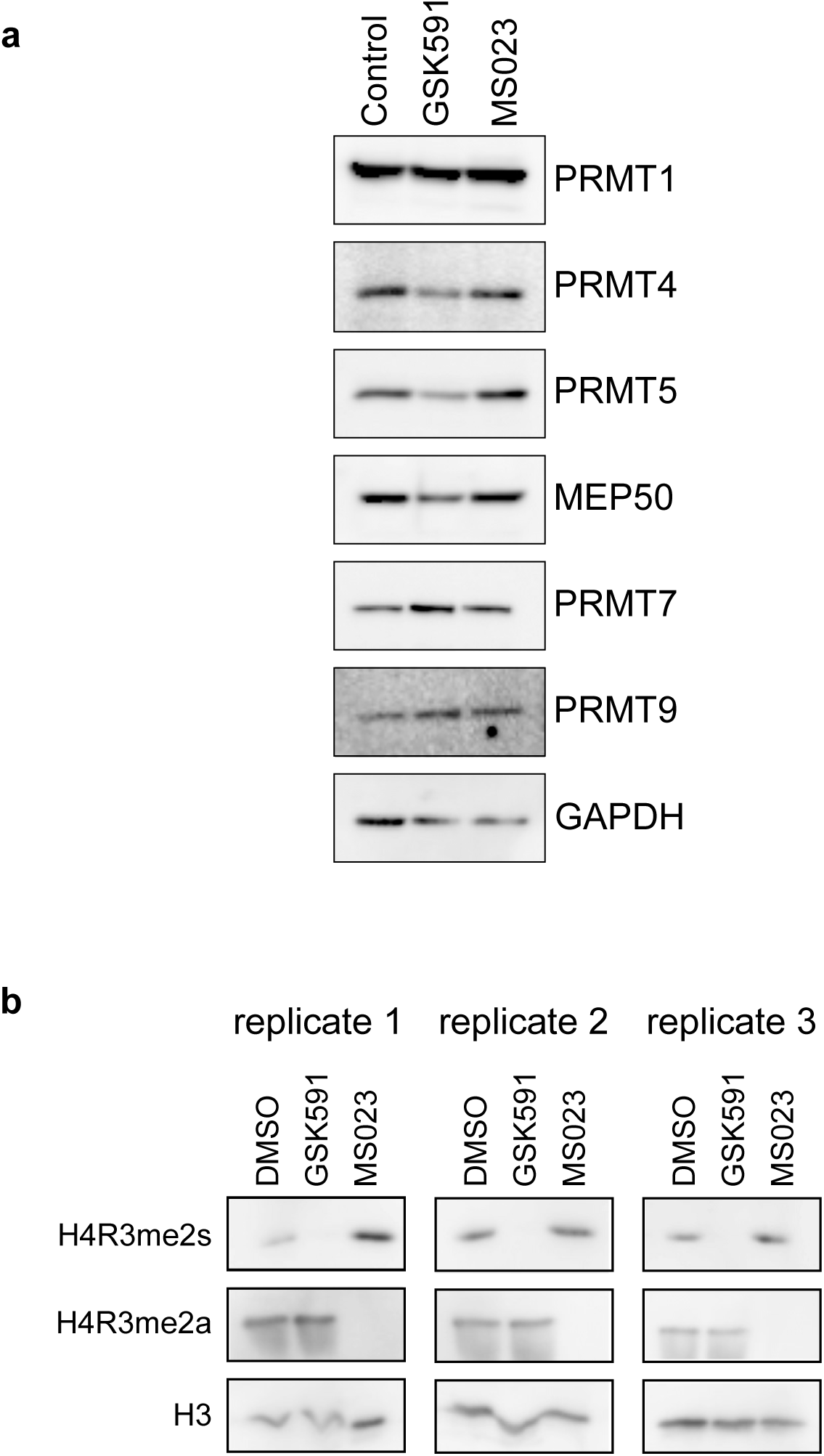
PRMT inhibition by GSK591 or MS023. **a.** A549 cells treated for 1 week with DMSO (control), 1 µM GSK591, or 1 µM MS023 were blotted for PRMT1, PRMT4 (CARM1), PRMT5, MEP50 (PRMT5 substrate presenter), PRMT7, PRMT9 and GAPDH (control). **b.** Histones extracted from three replicates of treated A549 cells, used for the subsequent PTMScan studies, were blotted for the PRMT5 catalyzed H4R3me2s and the type I PRMT catalyzed H4R3me2a. H3 was a loading control.

**Supplemental Figure S2.**
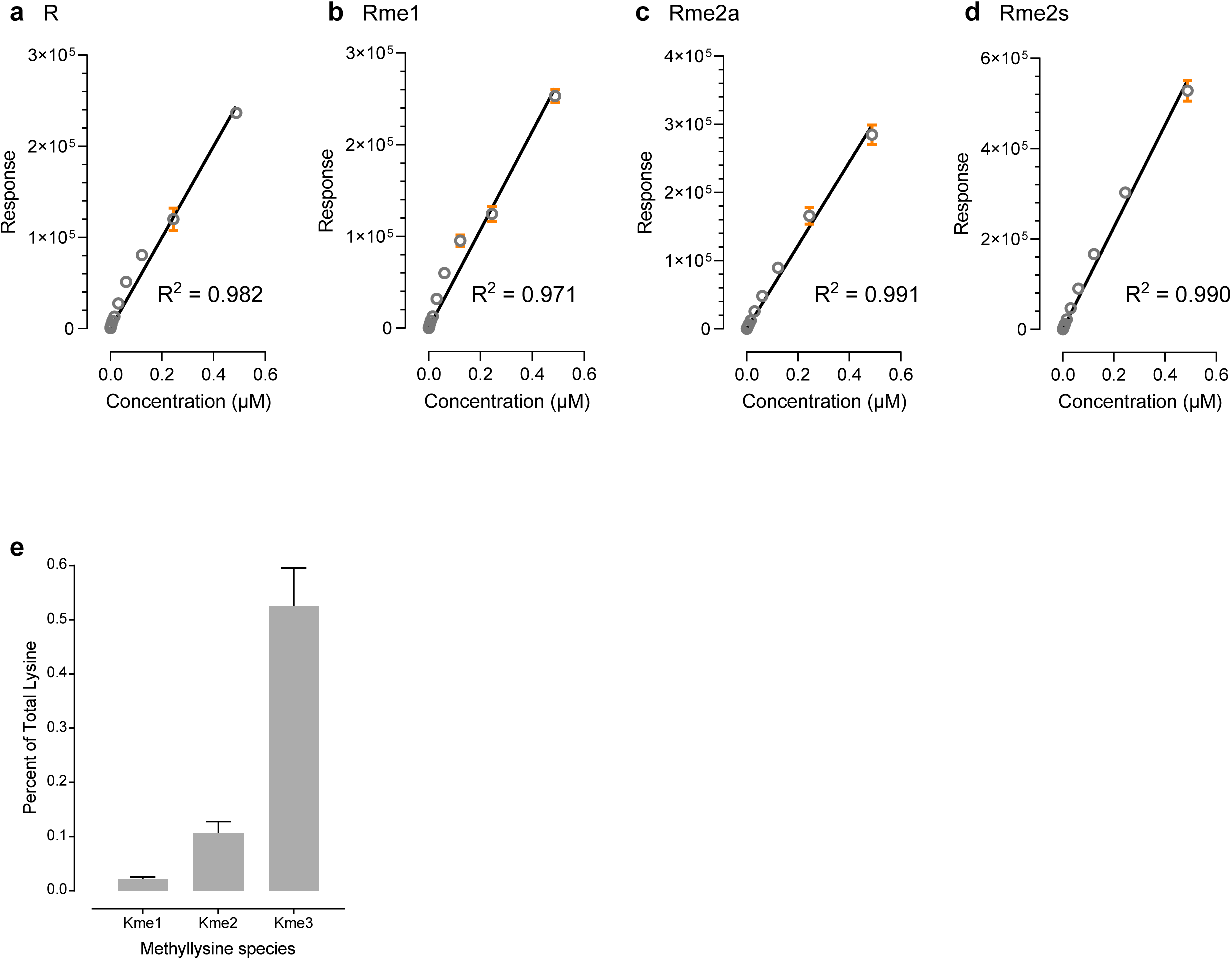
Amino acid analysis standard curve. **a.** Arginine response was measured, in triplicate, and plotted against known concentration (concentration determined by ^1^H-NMR with an adenosine standard). Replicates are shown; 95% confidence intervals are in orange. **b.** Monomethylarginine (Rme1) response measured in triplicate, as above **c.** Symmetric dimethylarginine (Rme2s) response measured in triplicate, as above **d.** Asymmetric dimethylarginine (Rme2a) response measured in triplicate, as above **e.** Calculated abundance of each methyllysine species as a percent of total lysine are shown

**Supplemental Figure S3.**
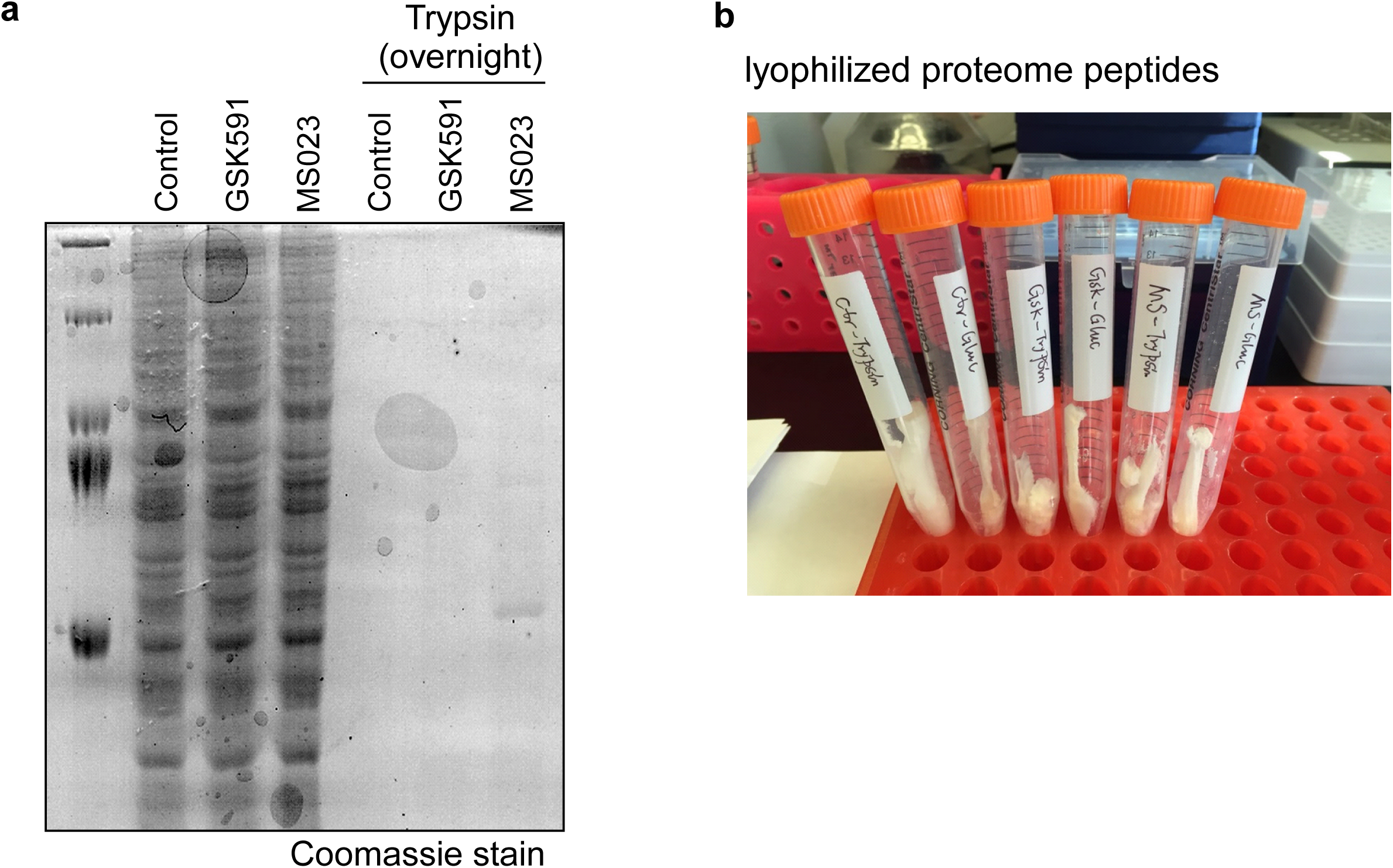
Proteome digestion for PTMScan. **a.** Coomassie stained gel showing the total proteome (left) and residual undigested protein after overnight trypsin incubation (right). All three conditions (DMSO control, GSK591, and MS023) are shown **b.** Photograph showing lyophilized tryptic peptides from each condition

**Supplemental Figure S4.**
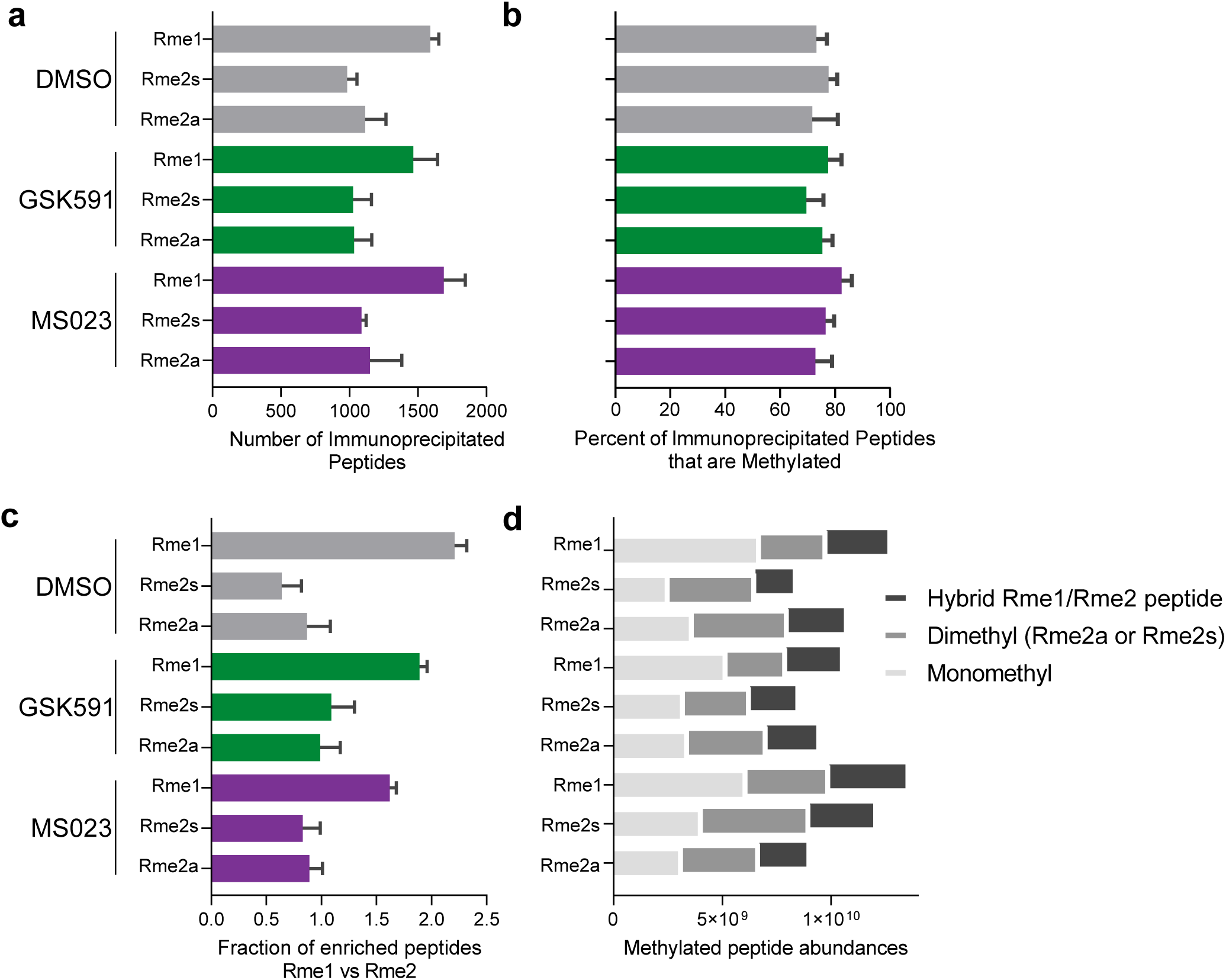
PTMScan peptide immunoprecipitation statistics. **a.** The number of peptides enriched in each of the three replicate sequential immunoprecipitations (Rme1, Rme2a, Rme2s) in each condition (DMSO in gray, GSK591 in green, and MS023 in purple) are shown. Error bars represent standard deviation. **b.** The percent of enriched peptides containing methylation in each of the three replicate sequential immunoprecipitations (Rme1, Rme2a, Rme2s) in each condition (DMSO in gray, GSK591 in green, and MS023 in purple) are shown. Error bars represent standard deviation **c.** The fraction of peptides enriched peptides that contain monomethylarginine (Rme1) versus dimethylation (Rme2s) in each of the three replicate sequential immunoprecipitations (Rme1, Rme2a, Rme2s) in each condition (DMSO in gray, GSK591 in green, and MS023 in purple) are shown. Error bars represent standard deviation. **d.** The stacked histogram shows the abundances of hybrid peptides (containing both Rme1 and Rme2), only demethylated peptides, or only monomethylated peptides enriched in each of the three replicate sequential immunoprecipitations (Rme1, Rme2a, Rme2s) in each condition (DMSO top, GSK591 middle, and MS023 bottom) are shown.

**Supplemental Figure S5.**
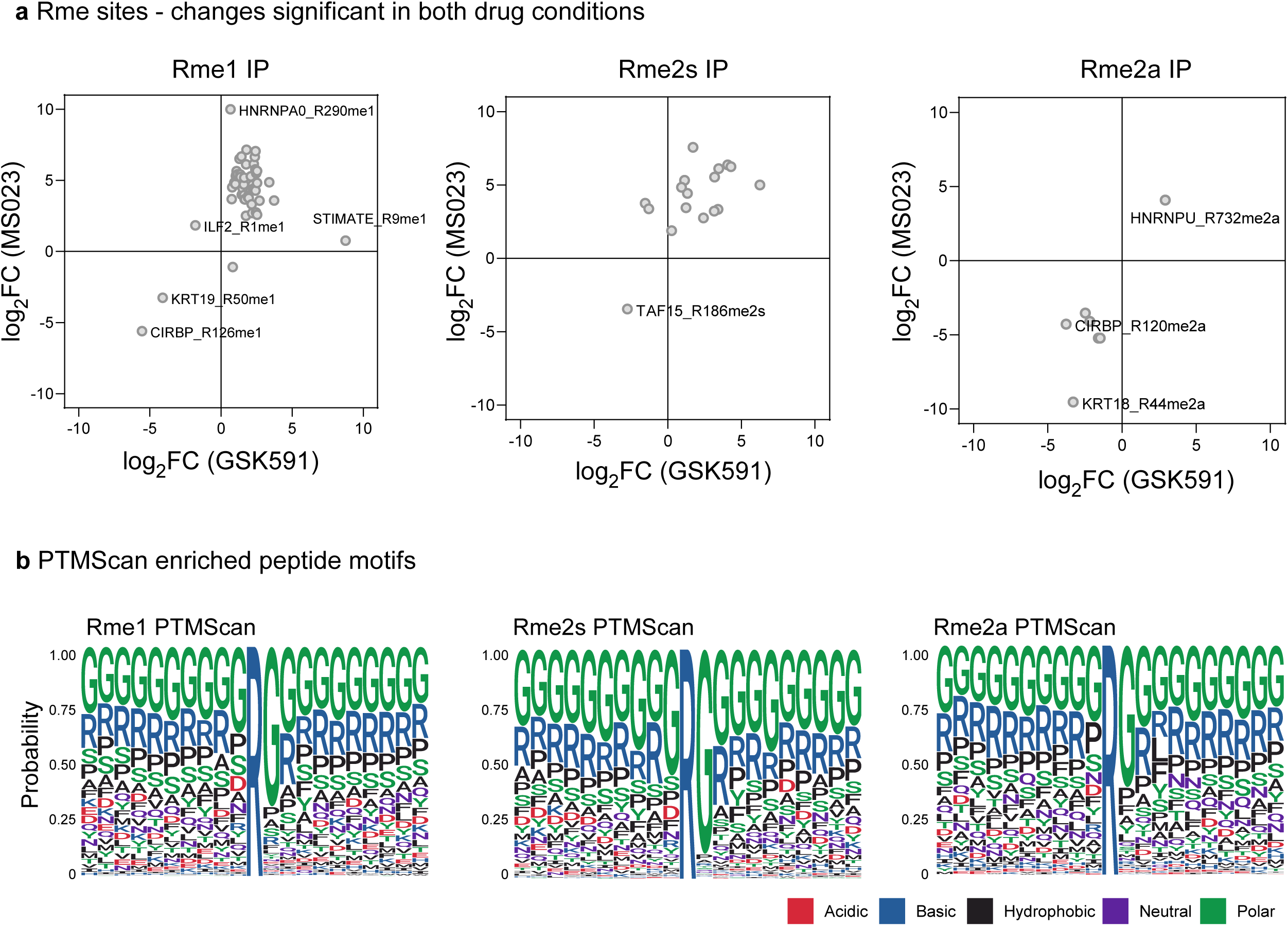
PTMScan enrichment characteristics. **a.** The log_2_(fold change) for methylation sites found in both the GSK591 (x-axis) and MS023 (y-axis) for monomethylarginine (Rme1, left), symmetric dimethylarginine (Rme2s, center), or asymmetric dimethylarginine (Rme2a, right) are shown **b.** Weblogo of Rme1, Rme2s, or Rme2a enriched peptides are shown. Residues shown span -10 to +10 amino acids from the methylated arginine R.

**Supplemental Figure S6.**
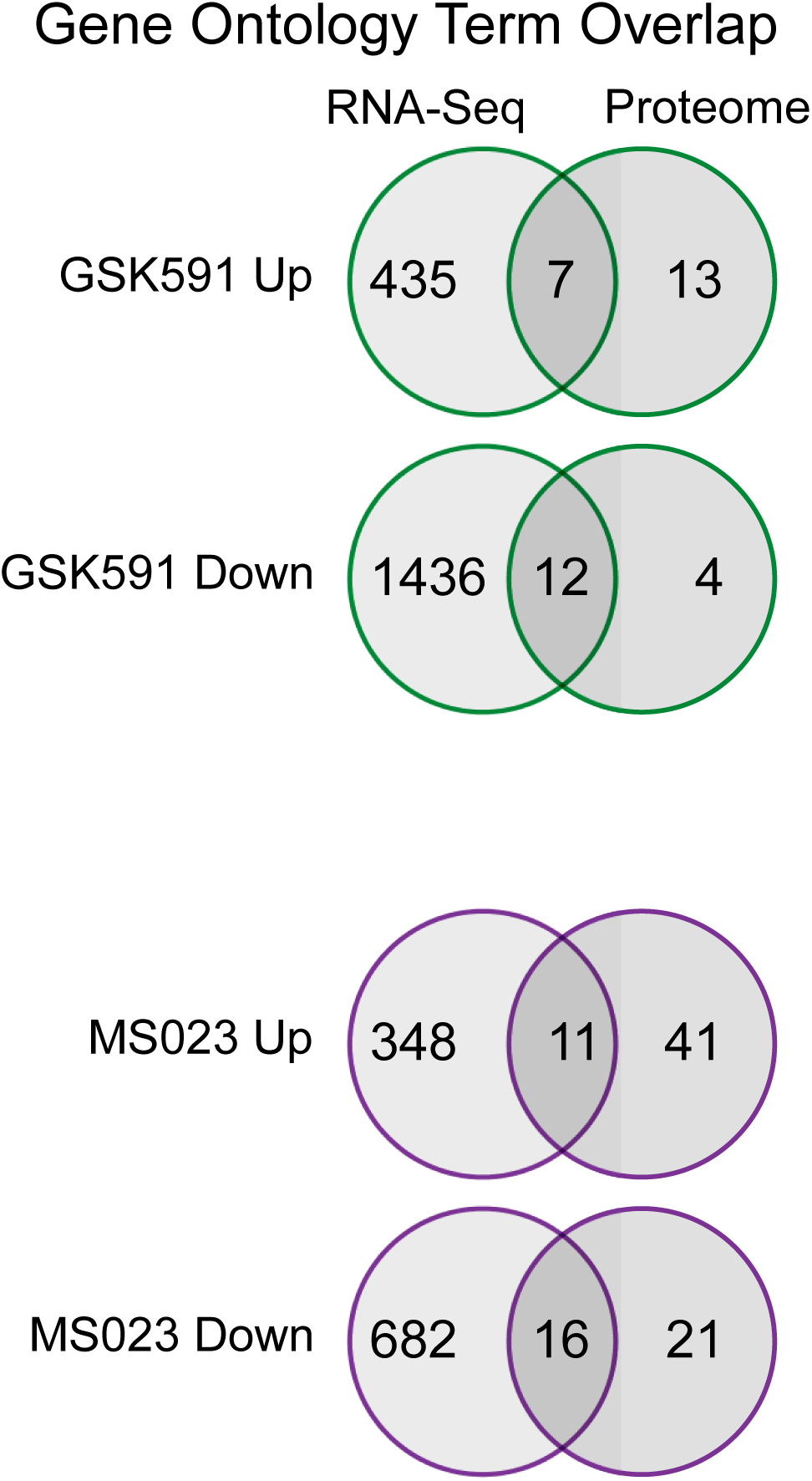
GO term overlap between RNA-Seq and Proteome datasets. **a.** Overlap of GO terms between up and down regulated genes (RNA-Seq) and proteins (Proteome) analyzed by gene ontology enrichment are shown. GSK591 is in green outline (top) while MS023 is in purple outline (bottom).

**Supplemental Figure S7.**
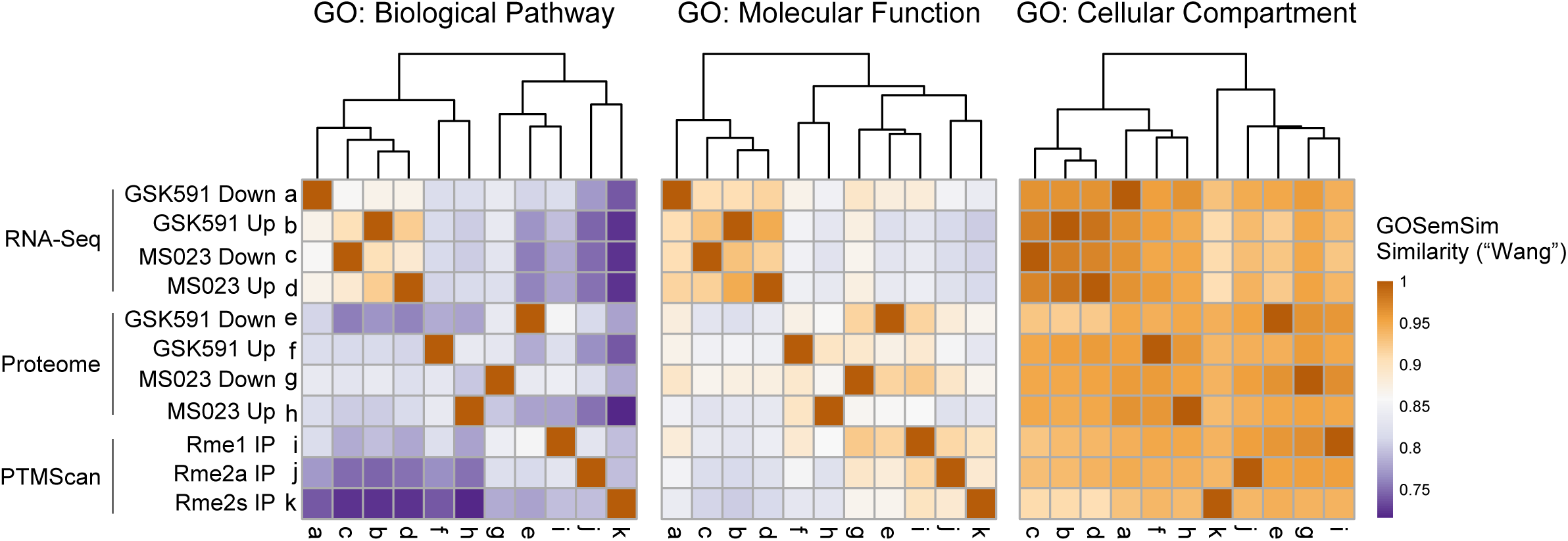
Summary of the results reveals that PTMScan only captures a subset of the cellular regulation by the family of PRMTs. GOSemSim matrices of gene ontology comparisons between the RNA-Seq, proteomic, and PTMScan altered genes and proteins. Biological pathway, Molecular Function, and Cellular Compartment ontology similarities, using the “Wang” semantic comparison metric, are shown as a heatmap.

***Supplemental Sheet 1 – PTMScan identified arginine methylation sites***

Shown are identified proteins (Uniprot Accession, Description, Gene name), their methylarginine site, whether or not the site lies in an intrinsically disordered region, the motif surrounding the R (+/- 10 aa), the protein name concatenated with the site, and their log2foldchange, p-value, and relative abundance in GSK591-treated cells relative to control and in MS023 cells relative to control. These data are from the PTMscan Rme1, Rme2s, and Rme2a immunoprecipitations in each of three conditions (Control, GSK591, and MS023 treated cells and are arranged in tabbed sheets in the Excel file. The final three tabs have the overlapping sites found significantly altered in both drug conditions.

***Supplemental Sheet 2 –PTMScan immunoprecipitated protein condition z-scores***

Shown are identified proteins (Description, Gene name) and relative enrichment row z-scores for each replicate of the PTMscan Rme1, Rme2s, and Rme2a immunoprecipitations in each of three conditions (Control, GSK591, and MS023 treated cells)

***Supplemental Sheet 3 – Proteome characterization of PTMScan input proteins***

Shown are identified proteins (Uniprot Accession, Description, Gene name) and their log2foldchange, p-value, and relative abundance in GSK591-treated cells relative to control and in MS023 cells relative to control. These data are from the total proteome.

